# Shoring up the base: the development and regulation of cortical sclerenchyma in grass nodal roots

**DOI:** 10.1101/2024.01.25.577257

**Authors:** Ian W. McCahill, Bahman Khahani, Cassandra F. Probert, Eleah L. Flockhart, Logayn T. Abushal, Greg A. Gregory, Yu Zhang, Leo A. Baumgart, Ronan C. O’Malley, Samuel P. Hazen

**Affiliations:** Biology Department, University of Massachusetts, Amherst, MA 01003, USA; Plant Biology Graduate Program, University of Massachusetts, Amherst, MA 01003, USA; U.S. Department of Energy, Joint Genome Institute, Lawrence Berkeley National Laboratory, Berkeley, CA, USA; Department of Human Genetics, University of Chicago, Chicago, IL, USA

**Author notes:** Author for correspondence: Samuel P. Hazen. **One sentence summary** Shoot-root anchoring in grass relies on cortex cell wall thickening activation by mechanical forces, gibberellic acid signaling, and the SWN7 transcription factor.

## Abstract

Plants depend on the combined action of a shoot-root-soil system to maintain their anchorage to the soil. Mechanical failure of any component of this system results in lodging, a permanent and irreversible inability to maintain vertical orientation. Models of anchorage in grass crops identify the compressive strength of roots near the soil surface as key determinant of resistance to lodging. Indeed, studies of disparate grasses report a ring of thickened, sclerenchyma cells surrounding the root cortex, present only at the base of nodal roots. Here, in the investigation of the development and regulation of this agronomically important trait, we show that development of these cells is uncoupled from the maturation of other secondary cell wall-fortified cells, and that cortical sclerenchyma wall thickening is stimulated by mechanical forces transduced from the shoot to the root. We also show that exogenous application of gibberellic acid stimulates thickening of lignified cell types in the root, including cortical sclerenchyma, but is not sufficient to establish sclerenchyma identity in cortex cells. Leveraging the ability to manipulate cortex development via mechanical stimulus, we show that cortical sclerenchyma development alters root mechanical properties and improves resistance to lodging. We describe transcriptome changes associated with cortical sclerenchyma development under both ambient and mechanically stimulated conditions and identify SECONDARY WALL NAC7 as a putative regulator of mechanically responsive cortex cell wall development at the root base.

## INTRODUCTION

In both monocots and eudicots, development of the root system begins with the emergence of the primary root at germination. These seed-borne roots then increase their effective interface with the soil by developing lateral roots, which originate from pericycle tissue at the periphery of the stele. However, in grasses like maize, wheat, and barley, vegetative development produces a cluster of nodes at the base of the plant. In response to phytohormone and environmental signaling, these nodes produce new roots, called nodal or shoot-borne roots, which together make up the shoot-borne root system, a characteristic aspect of grass root system architecture (Chochois et al., 2015; Sebastian et al., 2016). Anatomically, both eudicot and monocot roots have a central cylinder composed primarily of pith cells called the stele, which surrounds the xylem and phloem. The stele is surrounded by the cortex, several cell files of thin-walled parenchyma cells, through which water and solutes are radially transported from the soil to the vasculature in the stele (Lux et al., 2004). The cortex is separated from the stele by a single layer of cells called the endodermis, which develop an apoplastic barrier called the casparian strip, forcing solutes which may have been traveling between cells in the apoplastic pathway of the cortex to pass through the plasma membrane of the endodermis before entering the vasculature (Barberon, 2017; Barbosa et al., 2019). The cortex is finally surrounded by two additional cell layers, the exodermis, an outer apoplastic barrier that forms under certain conditions, and the epidermis, the outermost layer from which root hairs originate (Zimmermann and Steudle, 1998; Marzec et al., 2014).

Recent work has identified a novel and agronomically important developmental process in the cortical tissue of grass nodal roots: the formation of multiseriate cortical sclerenchyma (MCS) (Schneider, 2022). Compacted subsoils can create barriers to root growth that severely impact agricultural productivity. However, the roots of some grass crop species can traverse these barrier layers by developing MCS, several files of specialized cortex cells around the perimeter of the root. Multiseriate means “many rows”, reflecting that relative to other cortex cells, the MCS region contains very small cells, arranged in a tightly packed matrix. These cortex cells develop a secondary cell wall, a layer of cellulose and hemicellulose that is rigidified by the phenolic polymer lignin. Development of MCS layers allows roots to penetrate soils that genotypes lacking this trait cannot (Schneider et al., 2021).

Lodging, the failure of a plant to maintain its upright orientation, is often catastrophic for the plant as it often wounds the plant, impedes its access to sunlight, and exposes it to unfavorable microenvironments closer to the soil surface (Li et al., 2022). Lodging is a potent problem for growers of cereal crops worldwide and can cause drastic losses of grain yield when it occurs. Loss of upright orientation caused by a failure of the root system to maintain plant anchorage is called root lodging, in contrast to stem lodging, which is driven by breakages of the stem (Gardiner et al., 2016; Piñera-Chavez et al., 2016; Lindsey et al., 2021; Hostetler et al., 2022; Li et al., 2022). However, anchorage is the product of a larger shoot-root-soil system.

The dynamics of mature grasses subjected to strong winds illustrates the combined action of this multipart system in determining the ability to maintain anchorage. At the shoot, wind pushes air over the stem and leaves, exerting a bending force that depends on aerodynamic drag and stem rigidity. Because the plant is fixed at the base of the shoot, lateral wind force is converted to a rotational force acting on the plant, initially centered just below the shoot (Ennos et al., 1993; Brune et al., 2018). For nodal roots on the windward side of the plant, rotation will place them under tension, while roots on the leeward side will be compressed. As the plant is rotated further, windward roots of grasses tend to have sufficient tensile strength to resist failure, causing the plant to turn about a windward hinge point. Leeward roots must then either resist compression without buckling, move through the soil, or move with the soil as the zone of root-reinforced soil shears away from the bulk soil around it (Stubbs et al., 2019). Of these outcomes, only compression without failure maintains plant anchorage. Thus, in models of grass root system anchorage, the most important determinants of anchorage are soil properties to resist shear and compression and the resistance of nodal roots to bending and compression (Ennos et al., 1993; Goodman and Ennos, 1999; Ennos, 2000; Stubbs et al., 2019).

Modeling studies of wheat and maize anchorage note the presence of a ring of thickened, lignified cells at the periphery of the root cortex. These cells are specific to nodal roots and found only at the base of the root, where models predict resistance to compression is most important (Ennos, 1991; Ennos et al., 1993). While these studies indicate that this basal cortical sclerenchyma tissue very likely functions to promote anchorage by improving root resistance to buckling, neither the ontogenesis of these cells, nor the genetic regulation that promotes their wall development have been directly interrogated. However, micrographs of the basal region of nodal roots found in other literature suggest that basal cortical sclerenchyma development is a shared trait across many grasses. Among millet species grown under control or waterlogging conditions, *Panicum miliaceum*, *Setaria glauca*, and *Setaria italica* (but not *Panicum sumatrense*) developed similar lignified cortex cells at the root base, irrespective of treatment (Matsuura et al., 2022). Brace roots, a distinct class of nodal roots that originate above the soil surface in maize, had thickened cortex cells in the aerial portion of the root and conventional, parenchymatous cells below ground (Hoppe et al., 1986; Sparks, 2023). In an investigation of root development under drought, basal wheat cortical cells showed intense phloroglucinol-HCl staining around the root periphery. However, this lignified tissue was absent in the same region of rice nodal roots (Ouyang et al., 2020). Notably, while the presence or absence of MCS has not yet been interrogated in *Panicum* or *Setaria* species, MCS was present in wheat and maize, but absent in rice, suggesting a potential linkage between basal cortical sclerenchyma development at the soil surface, and MCS, which occurs closer to the root apex (Schneider et al., 2021).

For cortex cells to develop thickened, lignified cell walls in a specific region of the root requires that secondary wall biosynthetic processes are activated in some cells but not in others. Tissue and cell-type specific deposition of secondary walls in this manner is a common feature of other aspects of plant development. In general, the regulatory systems that control the biosynthesis of cellulose, hemicellulose, and lignin are well studied (Zhang et al., 2018; McCahill and Hazen, 2019; Coomey et al., 2020). In secondary cell wall depositing cells, a series of feed-forward transcriptional regulatory loops stimulate cellulose biosynthesis via the action of cellulose synthase complexes, hemicellulose biosynthesis via a number of glycosyltranferase families, and monolignol production via the phenylpropanoid metabolic pathway. In this study, we leverage the genetic model *Brachypodium distachyon* to interrogate the developmental regulation of basal cortical sclerenchyma in nodal roots, an understudied trait that is unique to grasses and has only been identified in crop species.

## RESULTS

### B. distachyon and other grasses develop a ring of cortex sclerenchyma in the basal region of nodal roots

We sampled leaf nodal roots of *B. distachyon* across a developmental gradient from the root apex to the shoot-root junction. These root segments were sectioned and stained with phloroglucinol-HCl, which produces an intense pink color in lignified tissues. To understand cortex cell wall thickening in relationship to the development of other root tissues, we quantified wall thickness and stain intensity of sclerenchyma, vascular, and apoplastic barrier cell types (**Supplementary Fig. 1**). The stele was smallest at the root apex and increased in size in older, more basal regions, both in absolute terms and as a percentage of the root cross sectional area (**Figs. 1A-C**). The walls of the pith cells that comprise the majority of the stele similarly increased in both thickness and stain intensity along this same gradient (**Figs. 1D,E**). Vascular cells within the stele likewise increased in stain intensity with tissue age, with outer metaxylem showing higher stain intensity than the much larger inner metaxylem. However, wall thickness of these cells did not vary appreciably along the root, indicating that these cells are fortified with lignin throughout development but do not undergo appreciable thickening. The apoplastic barrier tissues of the exodermis and the casparian bands showed a similar trend to xylem cells, with lighter staining at the apex and darker staining at the base, but relatively similar cell wall thickness throughout the root (**Figs. 1D,E**). Like the developing *B. distachyon* stem (Matos et al. 2013), we observed a positive trend with tissue age in both cell wall thickening and lignification for structural cells, while vascular cells lignified but did not significantly thicken as they developed.

**Figure 1.**
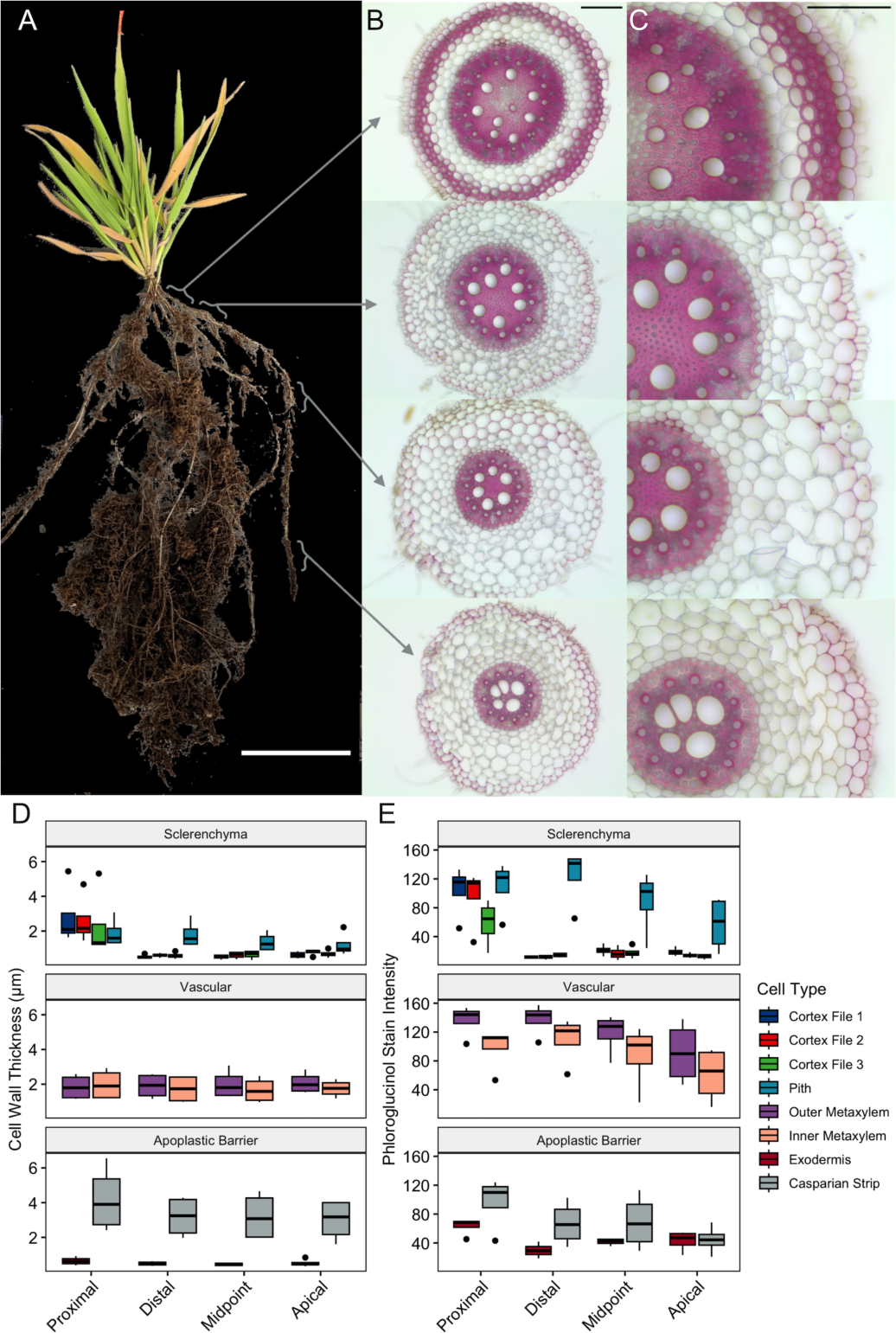
Histological characterization of *Brachypodium distachyon* nodal root development. Representative plant **(A).** Scale bar 5 cm. Phloroglucinol-HCI stained sections from the same root at 1Ox **(B)** and 20x **(C)** magnification. Scale bars 100 µm. Brackets and arrows indicate the root region corresponding to each section. Quantification of cell wall thickness **(D)** and phloroglucinol-HCI stain intensity (8-bit grayscale saturation value) **(E).** N = 4 plants. Plant means calculated from the mean of 15 measurements of cortex and exodermis, and casparian strip and 9 measurements for pith and metaxylem.

In contrast, cortex cell walls did not show a linear developmental trend in wall development. From the root apex to the distal basal region, cortex cell walls did not show appreciable phloroglucinol staining or secondary wall thickening (**Fig. 1**). However, exclusively in the proximal basal region, cortex walls in the outermost cell files were thicker and more intensely stained than any other cell type examined. These thick-walled cells had a morphology like the sclerenchyma cells found in mestome or interfascicular fiber tissues of the stem and formed a ring of heavily reinforced tissue around the root perimeter (**Figs. 1A-C**). Cortex cells were thickest in the first and second cell files (counting from the root exterior) and thinner but still distinctly thickened in the third cell file. While the other cell types examined showed global trends across the root, those cells were still similar between proximal and distal basal regions, indicating that those regions are of similar developmental age. Therefore, cortex wall thickening and lignification at the root base does not reflect a gradual process of cellular development and maturation, but instead might represent a specialized developmental program to reinforce the root where it emerges from the shoot.

To examine this trait directly in crop species, nodal roots from maize and wheat were examined. Because maize nodal roots have a region of red pigmentation near the shoot-root junction that resembles the pink color of phloroglucinol-HCl staining, the fluorescent stain Basic Fuchsin was used to characterize lignified tissues without confounding effects (**Fig. 2A,B**). Like *B. distachyon*, nodal roots of both species displayed thickened, lignified cortex cells, exclusively in the proximal basal region, giving way to thin-walled cortex cells in the distal basal region (**Fig. 2)**. In maize in particular, these thickened cells near the shoot-root junction resemble multiseriate cortical sclerenchyma, albeit with a lesser degree of cell wall thickening, and are similarly situated in the outermost cell files of the cortex (Schneider et al., 2021). This analysis confirmed that despite variations in scale and cortex anatomy between species, the lignification of cortex cells at the root base is conserved among both wild grasses and crops.

**Figure 2.**
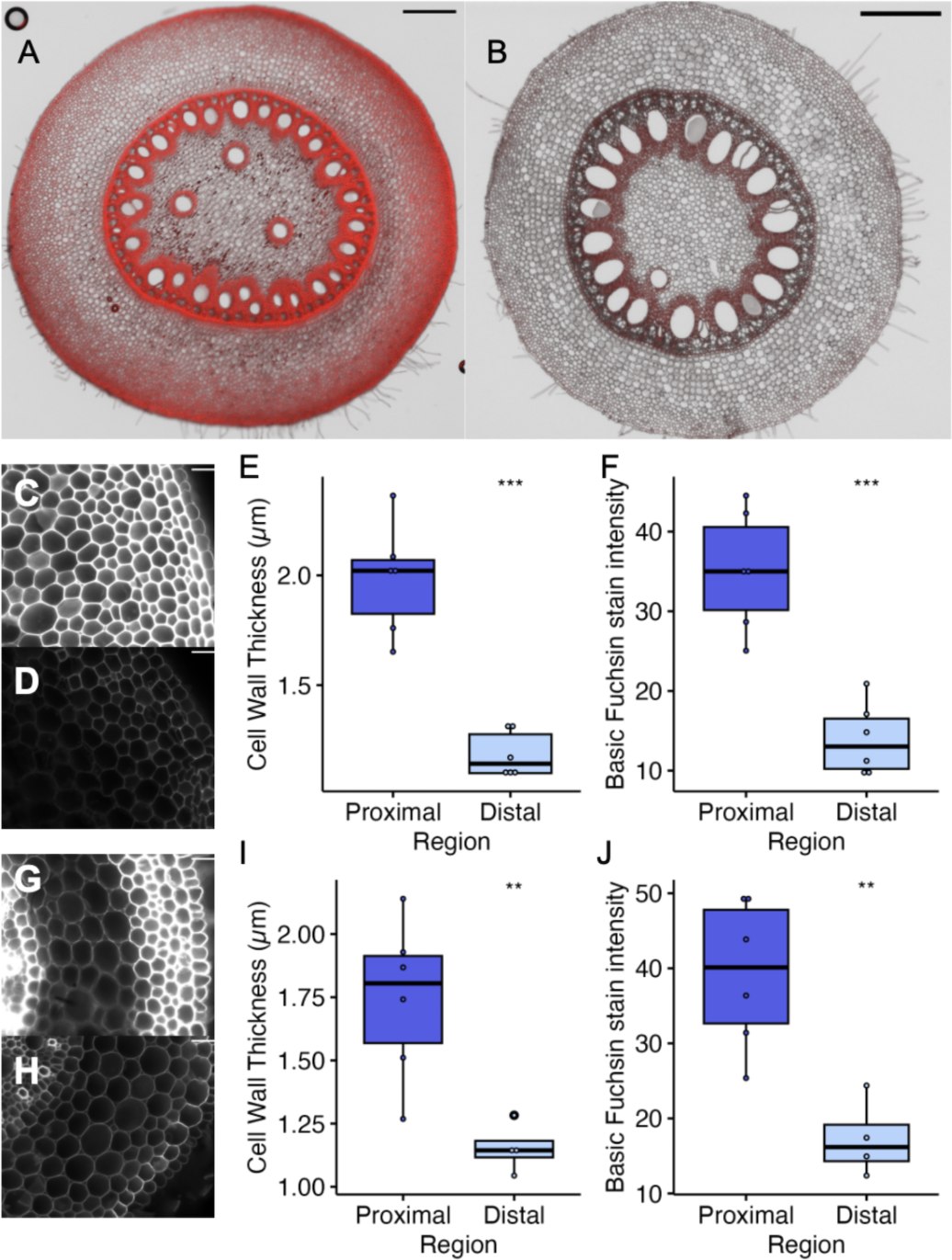
Identification of basal cortical sclerenchyma in crop species. Overlay of fluorescence (red) and brightfield (grayscale) micrographs of the proximal **(A)** and distal **(B)** basal regions of a Basic Fuchsin stained maize nodal root. Scale bars 500 µm. Characterization of secondary cell wall traits in maize **(C-F)** and wheat **(G-J).** Fluorescence micrograph of peripheral cortex cell in proximal **(C)** and distal **(D)** regions. Images shown in grayscale and brightened by the same factor for visualization. Scale bars 50 µm. Quantification of local cell wall thickness **(E,1)** and fluorescence signal **(F,J).** Student’s t-test, *\*\*p* 0.01, *\*\*\*p* 0.001,

### Cortex sclerenchyma develop continuously and undergo rapid cell wall thickening

To study secondary cell wall deposition specific to the root base, we examined development of this proximal tissue over time. As in the developmental gradient analysis, we observed different dynamics of secondary wall deposition based on cell function (**Fig. 3A-C**). Among both inner and outer metaxylem, stain intensity increased strongly over the course of the experiment, but cell wall thickness did not (**Fig. 3D,E**). At every time point, outer metaxylem cell walls were darker staining than inner metaxylem walls, although the magnitude of this difference decreased in later harvests. Exodermis cell walls likewise increased in stain intensity without a corresponding increase in wall thickness. However, these cells plateaued in stain intensity at 33 days after sowing (DAS). In the developmental gradient analysis, pith cells differed from other cell types in the degree of cell wall thickening. Here, these cells more than doubled in wall thickness, while increasing in stain intensity throughout the time series.

**Figure 3.**
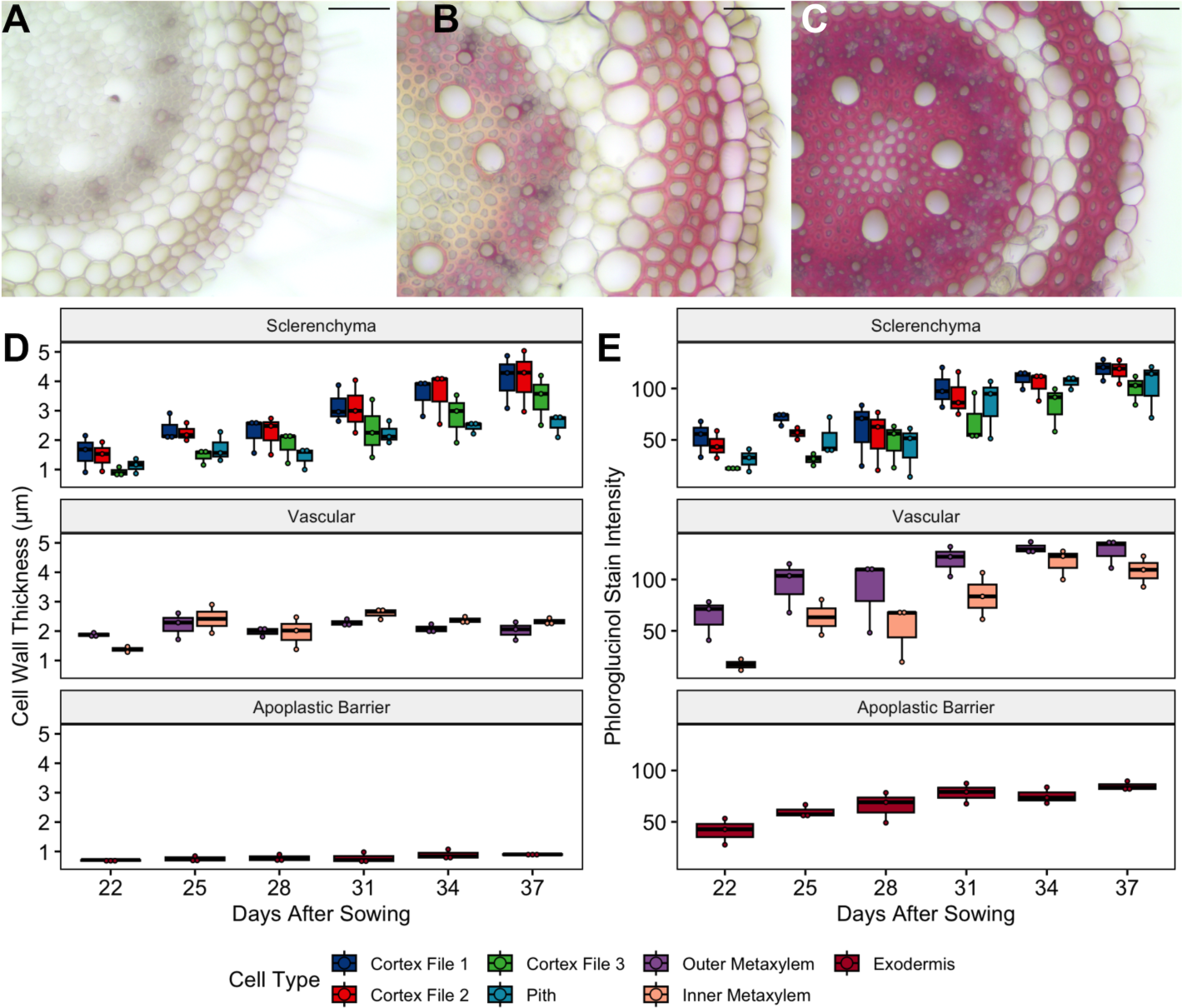
Time course of nodal root development. Phloroglucinol-HCI stained root sections, harvested at 22 **(A), 31 (B)** and 37 **(C)** days after sowing. Scale bars 50 µm. Quantification of nodal root secondary wall development. Cell wall thickness **(D)** and phloroglucinol-HCI stain intensity (8-bit grayscale saturation value)(E). Each point represents the cell type mean of one plant. Three roots of each plant were sectioned and 15 cells per root were measured for cortex and exodermis, 9 per root were measured cells for pith and metaxylem.

Cortex cells in the outer three cell files increased in both stain intensity and thickness throughout the course of the experiment. Notably, cell wall thickening and lignification first increased in the outermost cortex cells. In the earliest timepoint at which plants had developed nodal roots (22 DAS), cortex cell files 1 and 2 were already thicker and more intensely stained than pith cells. Cortex cells in the third cell files lagged the outer two files in wall development, only surpassing pith in thickness and staining at 28 DAS (**Fig. 3**). We also examined this data as a function of root length, irrespective of plant identity. This analysis showed similar patterns: that metaxylem and exodermis cells lignify without significant thickening, that pith and outer cortex cells thicken and lignify, and that cortex cell wall development begins in outer cell files (**Supplementary Fig. 2**). Thus, in terms of wall thickness, basal cortex cells develop faster than the baseline rate of sclerenchyma maturation but lignified at the same pace as other root cell types. Analysis of this data at the plant level suggests that across the shoot borne root system, basal cortex cells continuously thicken, even as other root cells reach maturity.

### Cortex sclerenchyma cells are lignified but not suberized

In many species, root cell layers that function as apoplastic barriers deposit the hydrophobic polymer suberin into their cell walls. Rice in particular develops a sclerenchymatous layer that is both lignified and suberized, that together with the epidermis, is called a complex hypodermis (Lux et al., 2004; Lynch et al., 2014). To assess whether thickened basal cortex cells in *B. distachyon* represent the development of a specialized multiseriate barrier layer or instead reflect altered development of otherwise-typical cortical cells, we separately stained basal sections from the same leaf nodal roots with phloroglucinol-HCl for lignin and Sudan Black B (SBB) for suberin. As expected of ligno-suberized apoplastic barrier tissues, exodermis cell walls and casparian bands were stained blue by SBB and pink by phloroglucinol-HCl (**Supplementary Fig. 3**). Outer cortex cell files, however, stained pink under phloroglucinol, but were not stained by SBB. That basal cortex sclerenchyma cells are lignified but not suberized suggests that these cells are functionally and developmentally distinct from the exodermis.

### Cortex cell wall thickening is stimulated by mechanical perturbation of the shoot

Models predict that grass root lodging resistance depends on the ability of the nodal root base to withstand the compression when the plant is displaced laterally by wind or other forces (Ennos, 1991; Ennos et al., 1993). To test whether these forces also stimulate the development of cortical sclerenchyma, we used an automated system to apply a bi-directional mechanical stimulus to plants at 90 min intervals. As in other mechanical treatment experiments, plants subjected to mechanical stress had significantly reduced stature (**Supplementary Fig. 4**) (Kouhen et al., 2023). Under ambient conditions, basal cortical sclerenchyma characteristically developed as a ring of thickened tissue around the root periphery (**Fig. 4A**). Mechanical treatment however, stimulated cell wall thickening and lignification throughout the cortex. To capture this global change, we quantified the local thickness and stain intensity of all cortex and stele cell walls. Touch treatment stimulated a significant increase in cortical cell wall thickness and Basic Fuchsin stain intensity but did not significantly affect these same measures of pith cells (**Fig. 4C-F**).

**Figure 4.**
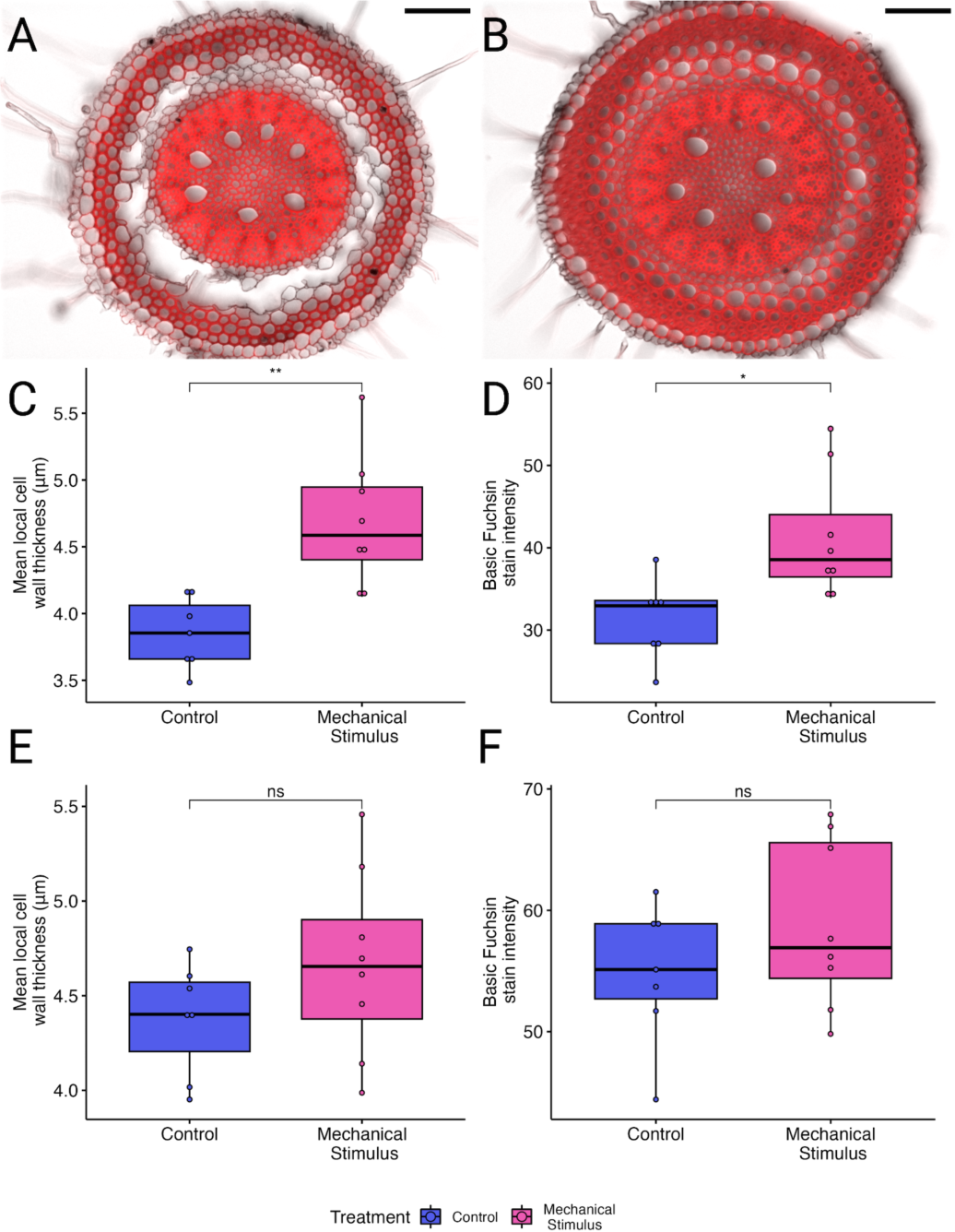
Mechanical stimulus induced wall thickening. Overlay of brightfield (greyscale) and Basic Fuchsin fluorescence (red) images of the proximal basal region of control **(A)** and mechanically stimulated **(B)** roots. Scale bar 100 µm. Quantification of cell wall characteristics. Wall thickness of proximal basal cortex **(C)** and pith **(E)** cells. Fluorescence intensity of proximal basal cortex **(D)** pith **(F)** cells. Tukey HSD pairwise comparisons, *\*p* s; 0.05, *\*\*p* s; 0.01, ns - not significant.

### Mechanically stimulated plants have increased resistance to artificial lodging

While the presence of stiffened roots at the shoot base is predicted to be a feature of the grass anchorage system, our system of mechanical treatment represents a novel avenue to manipulate the trait. To test whether increases in basal cortex thickening improve anchorage, we measured the force required to bend plants 30° from vertical (**Fig. 5A**). This approach is similar to other methods of assessing lodging resistance, which quantify the pushing or pulling forces required to rotate a plant to a given angle (Erndwein et al., 2020). Plants stimulated for 4 weeks required significantly more pulling force to deflect the stem, but only when tested perpendicular to the direction of treatment (**Fig. 5B)**. However, this may reflect that repeated treatment displaces soil away from the plant on the axis of touch, disrupting the root-soil interface required for anchorage.

**Figure 5.**
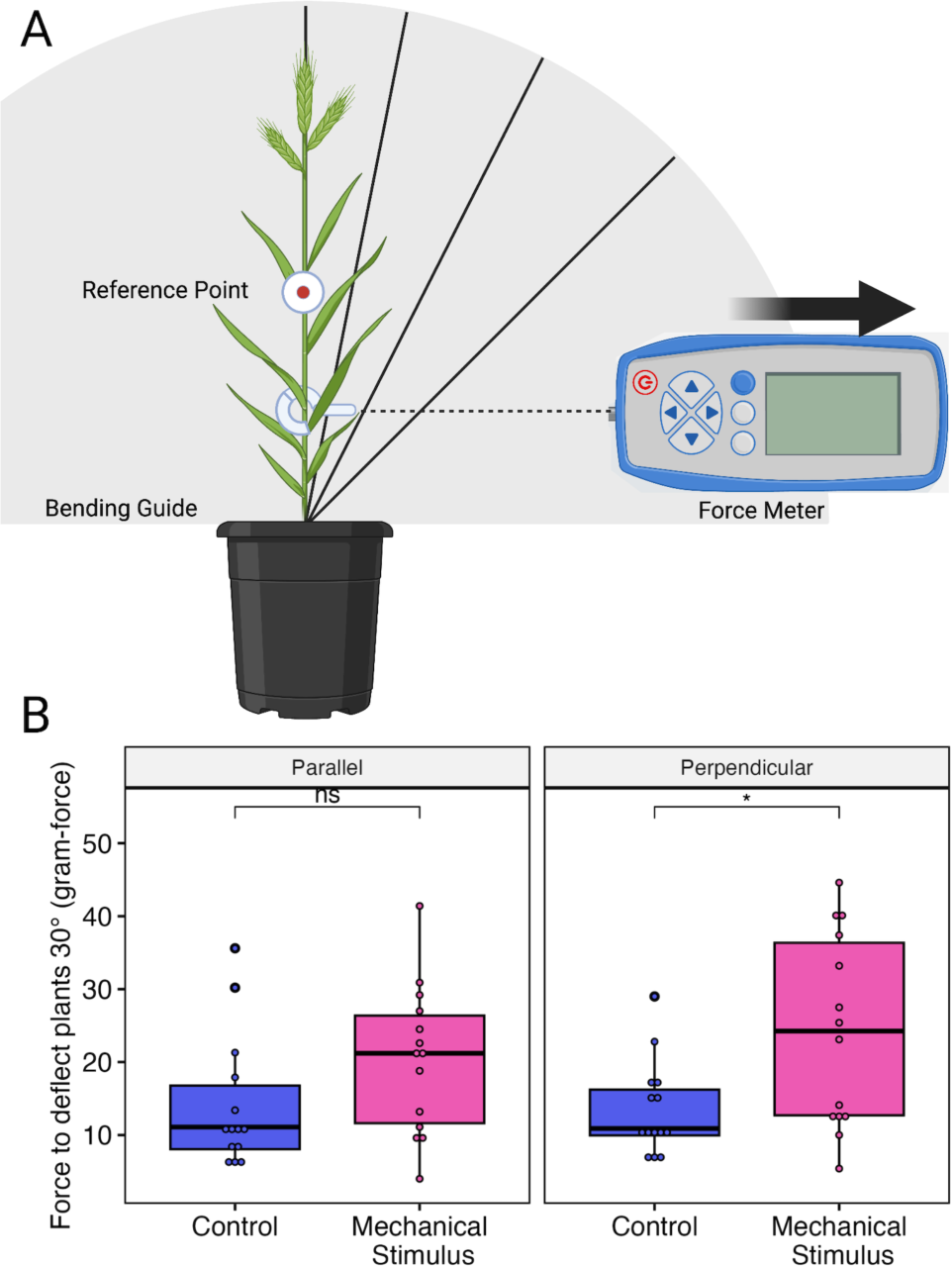
Testing resistance to lodging. Schematic of lodging resistance assay. **(A).** Peak force measured during 30° bending test **(B).** Tukey HSD pairwise comparisons, *\*p* 0.05, ns - not significant. N = 12-13 plants.

### Nodal roots increase their resistance to lateral compression in response to mechanical treatment

That mechanically stimulated plants have both thicker cortex cell walls and increased resistance to stem rotation is consistent with the hypothesis that nodal root stiffness allows grasses to maintain their anchorage. To more granularly evaluate how development under persistent challenge to anchorage shapes the material properties of individual roots, we measured the force required to compress different regions of single roots (**Fig. 6A**). Root diameter was similar across treatments and regions; as a whole, proximal and distal basal root segments did not have significantly different diameters, and treatment did not produce a difference within either tissue (**Fig. 6B**). In the distal basal region, control and stimulated roots were similar in the forces required to penetrate the root (**Fig. 6C**). Within control plants, the proximal basal region registered greater forces at the same relative penetration than the distal basal region, consistent with the hypothesis that cortex sclerenchyma improves the ability to withstand lateral compression. In proximal basal samples, roots of treated plants required more force to penetrate than control plants, further indicating that the additional thickening stimulated by mechanical perturbation provides additional resistance to mechanical challenge.

**Figure 6.**
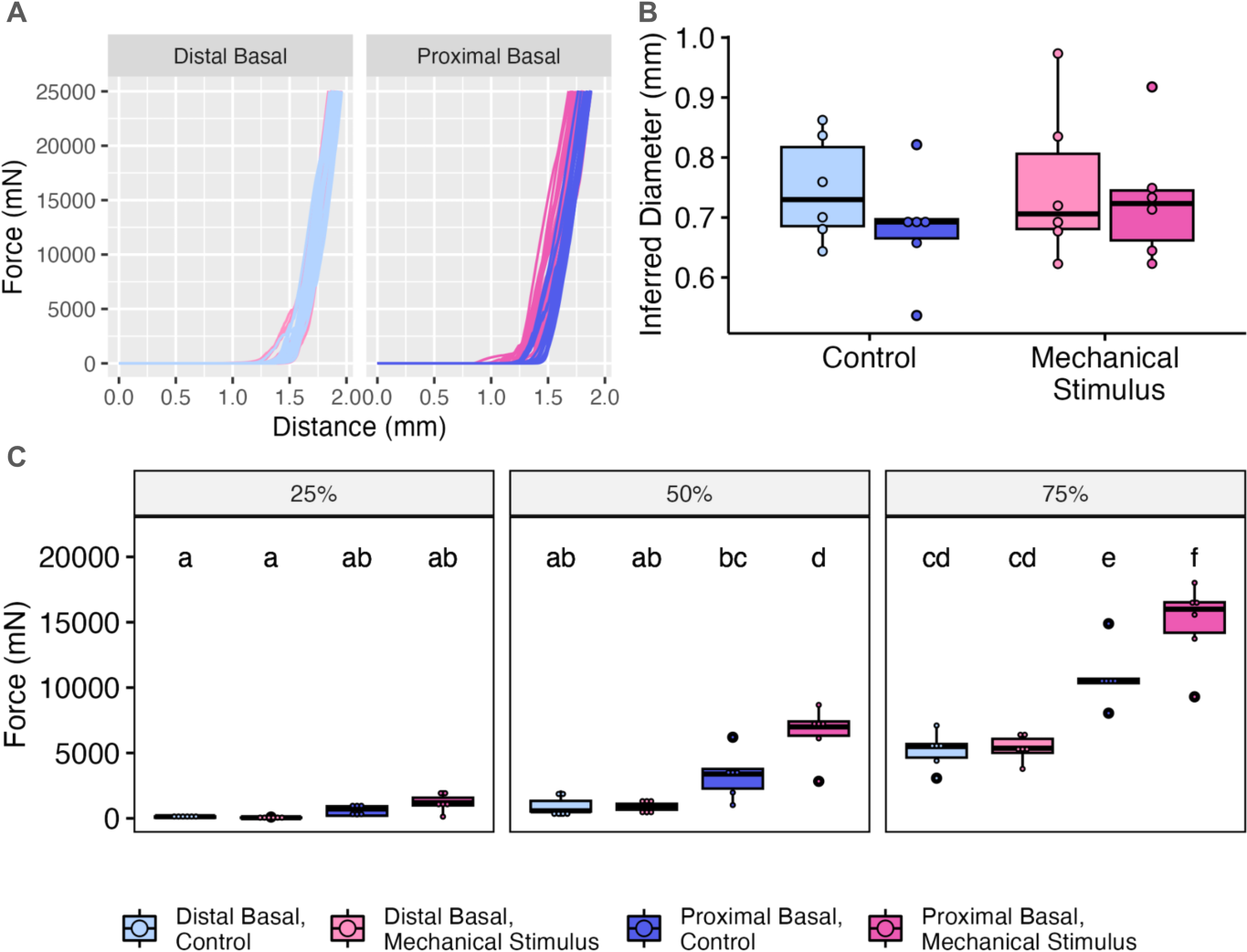
Testing root mechanical properties. Raw force-displacement curves **(A).** Root diameter inferred from testing data **(B).** Force measured at different percentages of root penetration **(C).** Each point represents mean force value for one plant, three roots per plant were assayed. N = 6 plants per treatment. Groups that do not share a letter are significantly different using Tukey’s HSD test, *p ≤* 0.05.

### Exogenous gibberellic acid stimulates root secondary cell wall development

Gibberellic acid (GA) phytohormone signaling is implicated in both secondary wall regulation and plant mechanoperception (Jiang et al., 2008; Wang et al., 2017; Singh et al., 2019; Coomey et al., 2023; Wu et al., 2023). To test whether GA is a component of the perception or transduction pathways that modulate cortical sclerenchyma thickening, we treated hydroponically grown plants with GA3, or the GA biosynthesis inhibitor, paclobutrazol. In general, nodal roots of hydroponically grown plants were markedly less lignified than those of similarly mature soil grown plants (**Fig. 1 and 7**). In contrast to mechanical treatment, which stimulated cortical sclerenchyma cell wall thickening but did not significantly affect the wall characteristics of pith cells, GA3 application dramatically increased stele phloroglucinol-HCl stain intensity, while also triggering wall thickening of the outer three cortical cell files **Fig. 7**). However, as with touch-induced cortical thickening, GA3 treatment did not trigger cortical cell wall thickening outside of the basal region where it is typically observed. Thus, while these results indicate that GA signaling promotes secondary wall thickening in cortical sclerenchyma cells as it does in other lignified cell types, they suggest that other regulatory mechanisms govern the initial transition between conventional cortex cells and cortical sclerenchyma (**Supplementary Fig. 5**).

**Figure 7.**
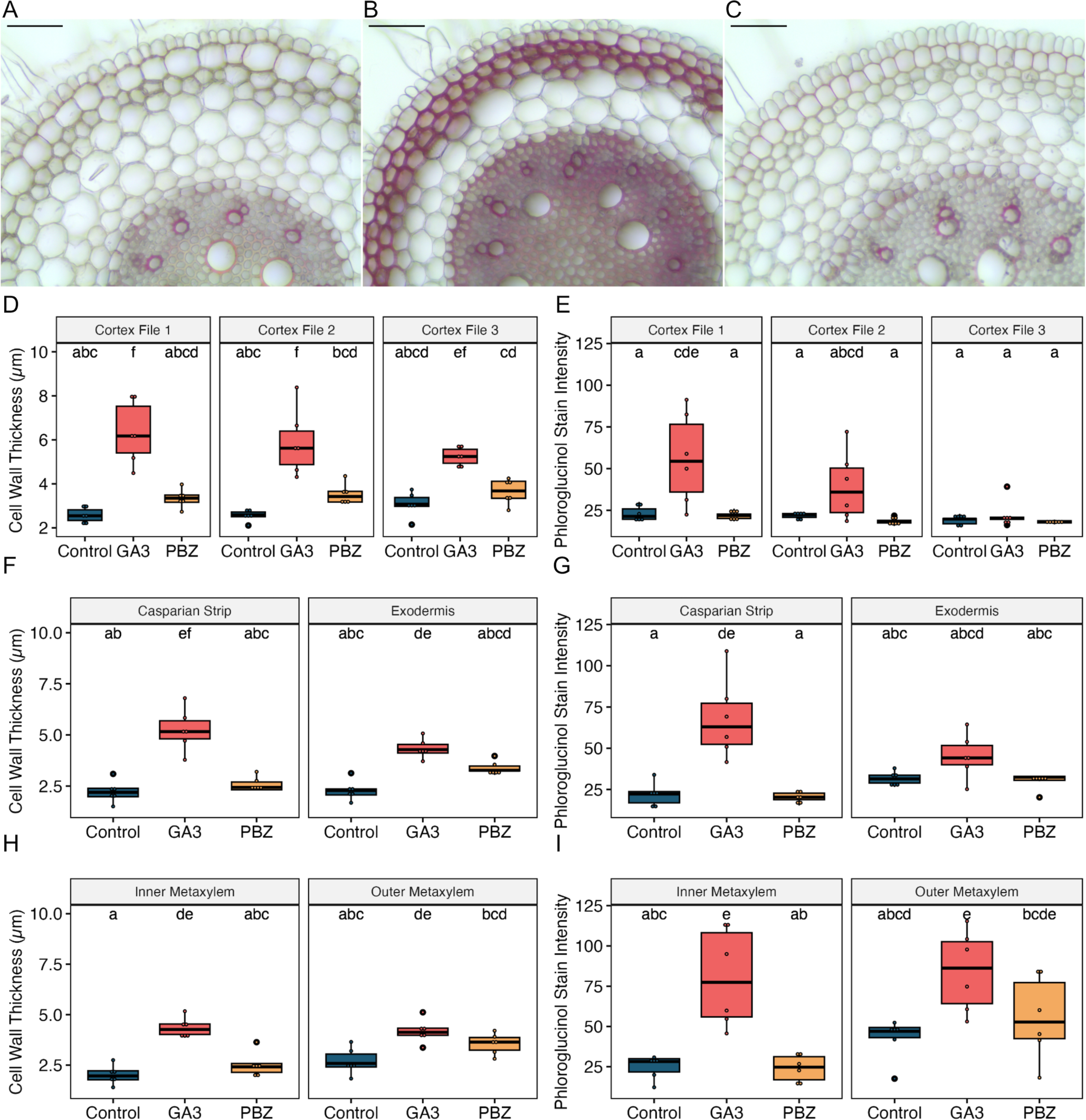
Phytohormone regulation of root secondary cell wall development. Transverse section of the nodal root base of plants grown under control **(A),** 10 µM GA3 **(B),** 0.1 µM paclobutrazol **(C)** treatment. Scale bars indicate 50 µm. Quantification cell wall thickness **(D,F,H)** and phloroglucinol stain intensity **(E,G,I).** Groups that do not share a letter are significantly different using Tukey’s HSD test, *p* ≤ 0.05.

### Transcriptomic signatures of basal cortex cell wall thickening

To uncover the transcriptomic signatures of cortical sclerenchyma development, we compared the transcriptomes of proximal and distal nodal root tissue (https://hazenlab.shinyapps.io/swiztc/). Further, to understand the genetic mechanism by which mechanical stimulus enhances cortical sclerenchyma thickening, we compared the proximal regions of control and mechanically treated plants. The transcriptome of proximal tissue samples, control and mechanically stimulated, were more like each other than the distal samples with more than three times as many genes differentially expressed between distal and proximal tissue of control plants than between control and treated proximal samples (**Supplementary Fig. 6**, **Supplementary Table 1-2**). Among the proximal and distal tissue differentially expressed genes, six cell wall related gene ontology (GO) terms were enriched (**Supplementary Fig. 7, Supplementary Table 3**). A notable 14 response to biotic and abiotic environment GO terms and lignin biosynthesis were enriched when comparing control mechanically stimulated proximal tissue.

To understand global trends in the root transcriptome across this set of treatment and tissue comparisons and the *cis*-regulatory elements that create them, we examined patterns of co-expression via hierarchical clustering analysis (**Supplementary Table 4, Supplementary Fig. 8**). Of the three largest clusters (1, 2, and 4), expression of genes in clusters 1 and 2 were positively associated with regions that undergo thickening, with cluster 1 showing stronger responses to mechanical stimulus, while cluster 2 genes showed similarly elevated expression in the proximal region of both control and mechanically stimulated roots, relative to distal untreated tissue. Genes in cluster 4 showed the opposite pattern and were most expressed in distal roots. Using a *de novo* k-mer approach to motif discovery, we next identified sequence motifs enriched in the *cis*-regulatory regions of genes in each cluster (**Supplementary Table 5**). Cluster 1, which contains the most distinct touch responsive expression pattern, showed significant enrichment for the Rapid Stress Response Element, P-box, G/A-box, AC element, and FAR1 binding sites, all of which have been implicated in transcriptional responses to mechanical forces (Walley et al., 2007; Van Moerkercke et al., 2019; Coomey et al., 2023). In Cluster 2, we also identified several distinct touch responsive elements including P-box, G-box, and site II elements which have been reported as a response to mechanical stimulus as well as several other motifs not associated with wall thickening. The Rapid Stress Response Element and W-box were significantly enriched in the Cluster 3 which have previously been described as touch responsive elements (Moore et al., 2022). The enrichment of this suite of motifs suggests that gene activation in response to mechanical stimulus is an important component of nodal root wall thickening.

Secondary cell wall deposition is in part governed by transcriptional regulation of genes in the various metabolic pathways that synthesize wall materials (Taylor-Teeples et al., 2015; Zhang et al., 2018; Coomey et al., 2020). The *B. distachyon* homologs of *Arabidopsis thaliana VASCULAR NAC DOMAIN* (*VND*) transcription factors that regulate vascular development, *SECONDARY WALL NAC* (*SWN*) gene (*SWN2*, *3*, *4*, *5*, and *6*) expression was extremely low and not affected by mechanical treatment (**Supplementary Fig. 9A**). *SWN7* and *SWN8* are most similar to *A. thaliana* activators of interfascicular fiber cell wall thickening, the *NAC SECONDARY THICKENING* (*NST*) and *SECONDARY NAC DOMAIN* (*SND*) genes (Valdivia et al., 2013). *SWN8* was more highly expressed in proximal tissue compared to distal tissue, but was not upregulated by mechanical treatment, while *SWN7* expression was both elevated in control proximal tissue relative to distal and mechanically stimulated. These NAC genes induce secondary wall thickening through direct regulation of wall biosynthetic genes, but also through activation of a larger number of MYB family transcription factors. Among the MYB genes with characterized roles in cell wall regulation, expression of *MYB2*, an ortholog of *AtMYB52/54* and *SWAM1*, which lacks an ortholog in the Brassicaceae, was elevated in proximal root tissues (Handakumbura et al., 2018). Consistent with a putative role in mediating touch-induced thickening downstream of *SWN7,* transcripts of *MYB104* and *MYB69* (orthologs of *AtMYB58/63* and *AtMYB20/42/85*, respectively) were elevated in proximal tissues, and much higher still in roots subjected to mechanical treatment (**Supplementary Fig. 9B**). Secondary cell wall cellulose is synthesized by the cellulose synthase genes *CESA4*, *7* and *8* and *CESA1*, *2*, *3*, *5*, *6*, and *9* are associated with primary wall cellulose biogenesis (Handakumbura et al., 2013). Here, no primary wall CESA gene was differentially expressed between tissues or treatments, while all three secondary wall CESAs were significantly upregulated in proximal tissue compared to distal (**Supplementary Fig. 10A**). Monolignols are synthesized through the phenylpropanoid pathway, and their polymerization into lignin is facilitated by several laccase enzymes. Comparing proximal and distal basal tissues, we measured increased expression of genes throughout the process of lignin biogenesis, including *PAL2, PTAL1, 4CL2, HCT1/2, CCoAOMT2, CCR1, F5H, COMT6, CAD1 and PMT* in the phenylpropanoid pathway and the laccases *LAC5* and *LAC8* (**Supplementary Fig. 10B**). In monocot cell walls, hemicelluloses, structural carbohydrates composed of a heterogeneous set of sugar monomers and linkages represent an especially large percentage of secondary wall biomass (Coomey et al., 2020). Analysis of known hemicellulose biosynthetic genes revealed no direct evidence of mechanical responsiveness, and only *GT47D3* was significantly upregulated in proximal tissues (**Supplementary Fig. 10C**).

To investigate SWN7 function as a putative transcriptional regulator of cortical sclerenchyma cell wall thickening, we characterized the genome-wide binding of SWN7 protein using DNA affinity purification sequencing (**Supplementary Table 6**). Analysis of the highest magnitude SWN7 binding peaks revealed a consensus SNBE/VNS element DNA binding motif characterized as the core binding motif for secondary wall regulating NAC transcription factors (**Fig. 8**)(Zhong et al., 2010; Nakano et al., 2015; Olins et al., 2018). Most of the binding peaks were in promoter regions or between the transcriptional start and stop sites.

**Figure 8.**
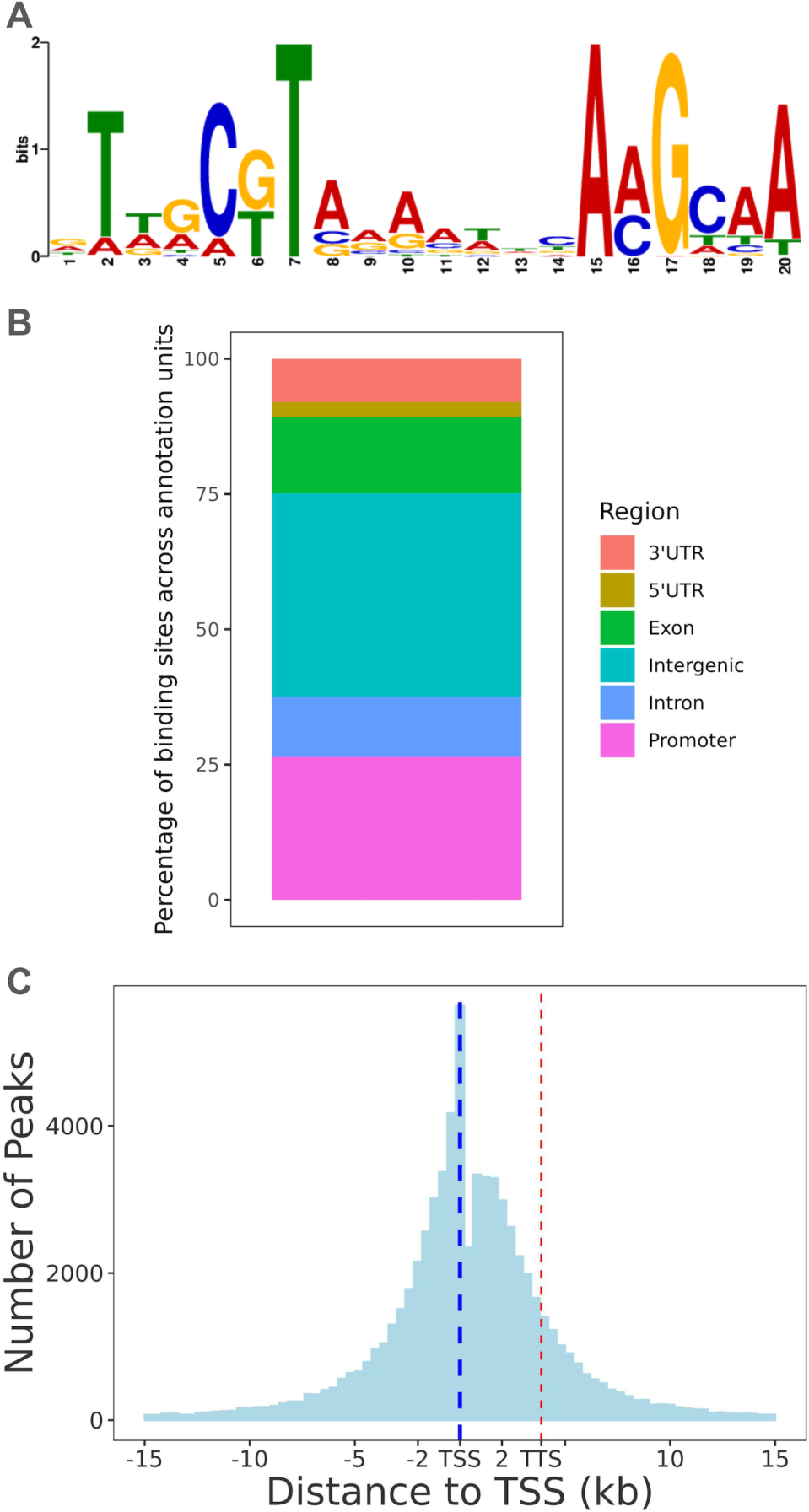
DNA affinity purification sequencing to determine SECONDARY WALL NAC7 binding sites. Most statistically enriched sequence motif in protein binding sites **(A).** Distribution of binding sites across genome annotation features, relative to primary transcripts of the *Brachypodium distachyon* annotation v 3.2 **(B).** Relative distribution of binding sites centered on the transcriptional start site {TSS, blue dashed line), transcriptional termination site (TTS, red dashed line) represents the average length of all annotated transcripts, approximately 4.5 kb away from the TSS **(C).**

Among the cell wall regulating MYB transcription factors, all the genes that showed differential expression by tissue or treatment also featured SWN7 binding peaks within the first exon or immediately upstream of the transcriptional start site (**Fig. 9, Supplementary Table 6**). This included *MYB104* and *MYB69*, which were strongly responsive to mechanical treatment as well as *SWAM1* and *MYB2*, which were only elevated in control proximal roots. Interestingly, the promoter of *SLR1*, which regulates GA biosynthesis, was bound by SWN7 and *SLR1* expression was significantly upregulated in proximal tissues (Huang et al. 2015). Considering cell wall carbohydrate biosynthesis, we found SWN7 binding peaks in the promoter regions of *CESA4, CESA8, COBL,* as well as in *GT47D3*, the only hemicellulose gene to show significant differences between root regions. Both *CESA4* and *GT47D3* had additional binding within the gene model itself, while CESA8 was notably bracketed upstream and downstream by two trios of binding peaks. SWN7 binding was associated with several lignin genes in different ways. *PAL2*, *F5H*, *HCT2*, and *COMT6* showed SWN7 binding both upstream and within an intron, and *PAL2* and *F5H* were bound 1 kb upstream of the transcriptional start site. We did not detect SWN7 binding upstream or in the gene body of *PTAL1* but did uncover a cluster of five significant SWN7 binding peaks downstream of the *PTAL1* locus. This aligns with the direct binding of SWN7 orthologs in various other species to the *cis*-regulatory region of cell wall biosynthesis genes and the MYBs that also regulate them (Zhong et al., 2008; Zhong et al., 2011; Huang et al., 2015; Taylor-Teeples et al., 2015; Zhong et al., 2015; Chen et al., 2019; Zhao et al., 2019).

**Figure 9.**
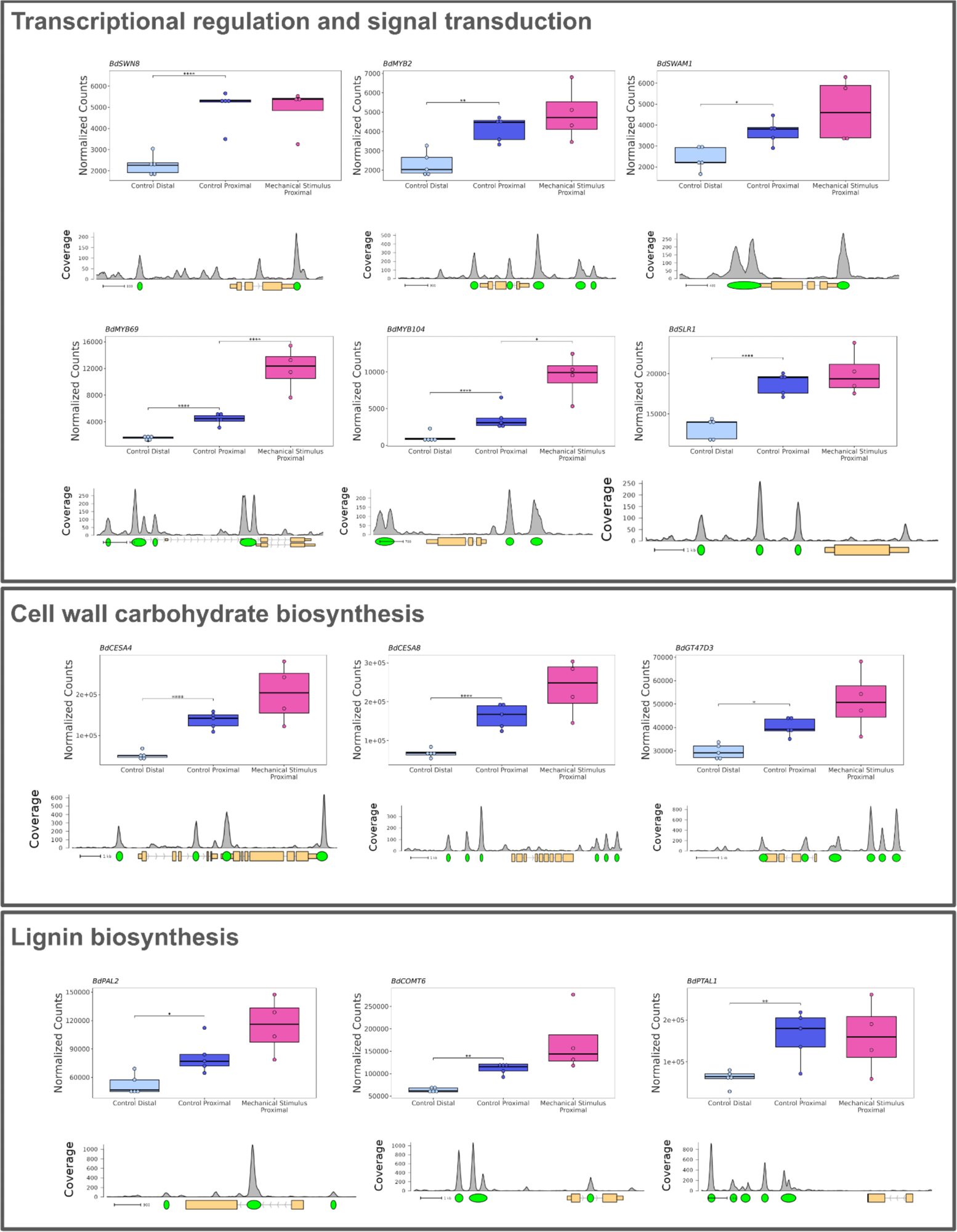
Secondary cell wall genes and the transcriptional regulators with expression associated with wall thickening in nodal roots were often DAP-seq binding targets of SECONDARY WALL NAC7 protein. Box plots are the average transcript abundance of 4-5 biological replicates for each tissue/treatment. Binding site determined as peaks of sequence alignment and denoted by green ovals. Scale bar unit is bases. Direction of transcription is shown with arrows on the gene model, 5’ and 3’ UTRs are depicted by narrowed rectangles on the gene model.* adj-p ≤ 0.05, ** adj-p ≤ 0.01, *** adj-p ≤ 0.001, **** adj-p ≤ 0.0001.

## DISCUSSION

The process by which plants detect the existence of touch or other mechanical stimulus is called mechanoperception, while altered growth or development in response to mechanical force is called thigmomorphogenesis (Jaffe, 1973; Bacete and Hamann, 2020). A global model of the mechanoreceptive systems that allow plants to undertake these responses has not yet been elucidated. However, several components of mechanosensing systems have been described. These include a suite of mechanosensitive ion channels that control calcium and other ion fluxes in response to membrane stretching (Kurusu et al., 2013; Monshausen and Haswell, 2013). Plants also detect immediate downstream effects of mechanical force. The receptor-like kinases FERONIA and THESEUS govern mechanical responses and maintain cell wall integrity by detecting carbohydrate molecules released from the wall by mechanical damage or other stress (Cheung and Wu, 2011; Shih et al., 2014; Bacete and Hamann, 2020).

Basal cortical sclerenchyma; cortex cells proximal to the shoot-root junction with thickened, lignified walls have been detected in several cereal crop species (Hoppe et al., 1986; Ouyang et al., 2020). Characterizing this trait in *B. distachyon*, we show that cortical sclerenchyma thickening is an output of a regulatory system by which grasses monitor mechanical stresses and proactively maintain their anchorage in the soil.

Across a suite of histological experiments, thickened cortex cells were universally present at the base of mature leaf nodal roots, but absent in more distal tissues, where roots are physically protected by soil (Ennos et al., 1993). In response to a mechanical treatment that transduced a bending force from the shoot to the root, cortex cells in the stressed region showed even stronger thickening. We conclude that basal cortical sclerenchyma development is stimulated by mechanical forces, with cortex thickening under unperturbed conditions reflecting the ambient forces that are inherent to supporting the plant’s mass.

Several instances of mechanically induced cell wall thickening have been previously described. In *B. distachyon*, a similar mechanical treatment induced strong cell wall thickening in interfascicular fibers, a type of sclerenchyma tissue in stems (Gladala-Kostarz et al., 2020). However, above-ground, mechanical treatment did not stimulate secondary wall development in cell types that usually lacked it, as we observed with cortical sclerenchyma. Furthermore, in our analysis of mechanically induced thickening in roots, we only observed responses in cortex cells, and not in pith sclerenchyma, suggesting that mechanoperception interacts with cell identity to control wall thickening under mechanical stress. In the woody dicot *Spartium junceum*, the lignin content of the root system increased when plants were grown on a slope (Scippa et al., 2006). Deliberate bending of *Populus nigra* taproots likewise altered root morphology and increased root lignin content (De Zio et al., 2016). While these responses indicate that dicots also use cell wall deposition to meet the mechanical demands of root anchorage, they are distinct from the tightly localized, cortex-specific thickening of grass nodal roots that we describe here.

The topography of the grass secondary cell wall network is well-established, with NAC transcription factors in the SWN family acting upstream of a more numerous group of MYB genes, while also directly activating wall biosynthesis (Coomey et al., 2020). Of NAC transcription factors with established roles in secondary wall regulation, SWN7 was the only gene in this group to show a significant transcriptional response to both ambient and strong mechanical forces. We therefore identify SWN7 as a putative upstream regulator of touch-activated cell wall thickening in roots. DAP-seq analysis of SWN7 protein binding further supported the hypothesis that cortical sclerenchyma development is a consequence of mechanical activation of SWN7. We confirmed that SWN7 binds the VNS-element and identified SWN7 binding peaks for *SWAM1*, *MYB2*, *MYB69* and *MYB104* as well as several biosynthetic genes, including *CESA4* and *CESA8*. Analyzing the promoter regions of genes that showed positive responses to ambient or strong forces (Clusters 1 and 2), we found enrichment of the MYB-associated binding site AAATATCT, as well as the AC-element, which is bound by SWAM1 (Handakumbura et al., 2018). Thus, our DAP-seq and RNA-seq experiments suggest a mechanism in which SWN7 is stimulated by touch and subsequently directs cortical wall thickening, both directly and through activation of MYB intermediaries.

In rice, the SWN7 ortholog OsNAC29/OsSWN2 is negatively regulated by a physical interaction with SLR1. Exogenous GA triggers SLR1 degradation, allowing OsNAC29 and other NAC genes to activate secondary wall thickening (Huang et al., 2015). We likewise observed strong activation of cell wall thickening under GA treatment, but found that unlike mechanical treatment, GA was not sufficient to stimulate cell wall thickening in inner cortical cell files. Therefore, it is likely that GA stimulates cell wall thickening in cortex cells by relieving post-translational repression of SWN7-directed thickening, but only in peripheral cortex cells that have already gained a sclerenchymatous cell identity through ambient mechanical forces. A model of ambient and strong mechanical effects on cortical sclerenchyma development is shown in **Fig. 10**.

**Figure 10.**
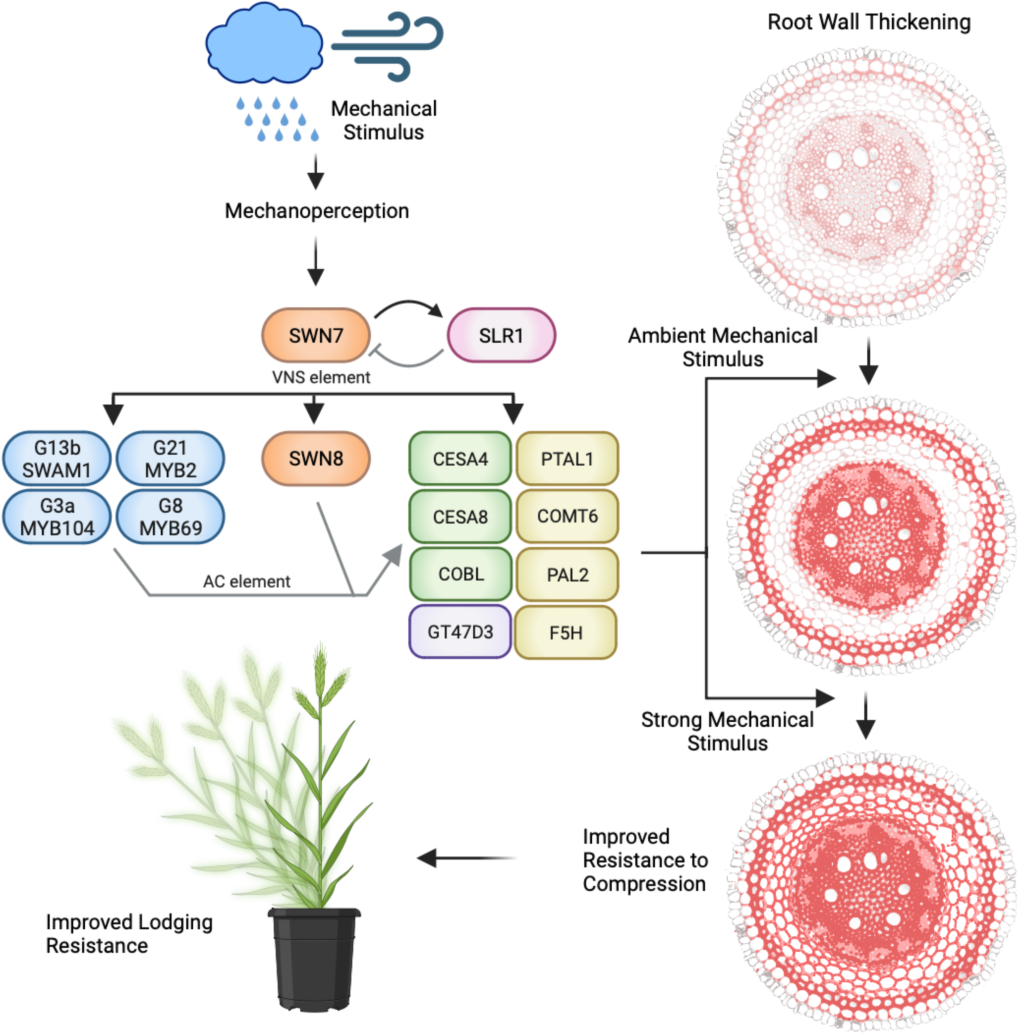
Model of the effect of mechanical stimulation on transcriptional regulation of nodal root basal cortex secondary wall thickening. NAC, GRAS, and MYB family DNA binding transcription factors are orange, pink, and blue ovals, respectively. Cellulose associated *CELLULOSE SYNTHASE* As and *COBRA-like (COBL),* green rectangles, hemicellulose associated *GT4303,* violet rectangle, and lignin, yellow rectangles, genes are binding targets of the transcription factor proteins. Arrows indicate activation and bars repression. Black lines are supported by RNA­ and OAP-seq results in this study. Connections depicted by gray lines are previously described in the literature for various species (reviewed by Rao and Dixon, 2018; Zhang et al., 2018). Mechanical forces acting on the stem induce bending at the base of the root and cells at the root periphery begin to thicken. Under ambient conditions, thickening of outer cell files stiffen the root against bending, protecting interior cortex cells. Under strong mechanical stimulus inner cell files develop thick walls.

Cell wall remodeling governed by this process is essential to allow proper growth and prevent catastrophic mechanical failure of exterior cells. In *B. distachyon* roots, certain cortex cells undergo thigmomorphogenic cell wall thickening around the root exterior. This process alters the way roots interact with additional mechanical forces, and presumably affects cell-cell interactions between interior cortex cells and thickened cortex sclerenchyma. Elucidation of the mechanisms that govern the perception of these mechanical stimuli, and that ensure that thigmorphogenically stimulated roots maintain their viability will further clarify the mechanisms by which grasses proactively maintain their anchorage and resist lodging.

In characterizing basal cortical sclerenchyma, we note the visual similarity between that trait and multiseriate cortical sclerenchyma, particularly in our analysis of maize. In plants that develop MCS, thickening of cortex cells deeper in the soil allows roots to resist failure as they penetrate compact soils (Schneider, 2022). While MCS has been detected in wheat, maize, and barley, it is absent in landraces of those species, and only found in more modern cultivars (Schneider et al., 2021). This apparent pattern of parallel evolution strongly suggests that, in selecting for yield in environments with some degree of soil compaction, breeders inadvertently selected for mutations that exaggerated that activity of a cortex-thickening pathway present in grasses. In describing the mechanical and genetic regulation of cortex thickening in *B. distachyon*, a wild relative of these species, this work may provide a platform to better understand the evolution of an important agronomic trait.

## Materials and Methods

### Plant growth conditions

*B. distachyon* accession Bd21-3 seeds were imbibed for 10 days at 4 ℃ then planted in potting mix (Pro-Mix BX Mycorrhizae) in 22 cm deep Conetainer pots (Ray Leach SC10). Plants were grown in a PGC-105 growth chamber (Percival Scientific) with a ∼450 μmol·m^−2^·s^−1^ day period of 16 h at 24℃ and a night period of 8 h at 18℃. Fertilization with Grow 7-9-5 fertilizer (DYNA-GRO) diluted to 0.6 ml/liter began ∼2 weeks after germination. For the time course analysis, fertilization was performed once a week via bottom watering. For all other growth chamber experiments, fertilization was delivered via top-watering every other week. In the time-course experiment, plants were harvested at 3-day intervals after the onset of nodal root development, and the longest 3 roots were selected. For all histological and mechanical characterization of the root base, we define the proximal basal region as the interval from 0-2 cm, and the distal basal region as the interval from 4-6 cm, with 0 representing the shoot-root junction. Maize inbred B73 and wheat cultivar Kronos were grown as described above. The mechanical stimulus apparatus used to make contact every 90 min, is previously described (Coomey et al., 2023). In the *B. distachyon* mechanical stimulus experiment, plants were grown for 2 weeks prior to treatment onset. Samples were harvested after 2 weeks of treatment for RNA sequencing, and after 4 weeks of treatment for histological and mechanical analysis.

For the hydroponics experiment, seeds were germinated on half strength MS plates, and then transferred to hydroponic growth in half-strength Hoagland’s No. 2 media (Caisson Labs). Aqueous solutions of paclobutrazol and GA3 were freshly prepared and diluted to 0.1 µM and 10 µM respectively in hydroponic media. To allow for recovery after transfer, pharmacological treatment was applied after one week, and plants were harvested for histological characterization after two weeks of growth in control or treated media. To prevent stagnation or phytohormone depletion, the media was changed weekly.

### Histological characterization and image analysis

Root segments were embedded in 5.4% agarose and 60 micron transverse sections were prepared using a VT1000 S vibratome (Leica) with the following settings: Speed: 5;50, Frequency: 8, Feed: 60. Operating the vibratome in continuous mode, we sectioned the entire embedded region of each root. Phloroglucinol-HCl staining was performed according to (Liljegren, 2010). Sudan Black B staining was performed as described in (Ruzin, 1999). Basic Fuchsin staining was performed as described by (Kapp et al., 2015). Brightfield imaging of phloroglucinol-HCl and SBB sections was performed on an Eclipse E200 microscope (Nikon) and Pixelink camera (Pixelink). Phloroglucinol-HCl staining intensity was quantified using a saturation intensity approach (Lee et al., 2019). For cortex, pith, and exodermis cells, five unique cells were measured, while for metaxylem, at least two cells per image were measured. For each root, images of three separate sections were analyzed as above. Fluorescence and brightfield imaging of Basic Fuchsin stained sections was performed using a BZ-X800 fluorescent microscope (Keyence). Fluorescent images were captured with an excitation wavelength of 545 nm and an emission wavelength of 605 nm. In this experiment, five section images per root replicate were analyzed, and local cell wall thickness and stain intensity were quantified using a MIPAR based image segmentation algorithm (Sosa et al., 2014). In the hydroponic experiment, thickness and phloroglucinol intensity of cortical, xylem and apoplastic barrier cells were quantified as described above, while phloroglucinol-HCl staining of the stele was measured using MIPAR.

### Physical characterization of root anchorage

To assess resistance to lateral deflection, the force required to bend plants 30° from vertical was measured. Because *B. distachyon* developed more rigid stems in response to mechanical treatment in other studies, bending force was applied as close to the soil surface as possible to avoid the influence of stem characteristics (Gladala-Kostarz et al., 2020). Plants were placed in front of a bending guide with 10°, 20°, and 30° of deflection. A small sticker was applied to the tallest stem to serve as a reference point. To allow the plant to move, but minimize the influence of stem elasticity, a flexible metal hook, constructed from 18-gauge aluminum wire, was secured around the base of each plant, ∼2 cm above the lip of the conetainer. The hook was connected by a ∼15 cm length of sewing thread to a DS2-1 force gauge (IMADA), set to peak mode. The force gauge was gently pulled along a guide surface, in line with the plant, until the stem came in line with the 30° guideline. Two measurements were performed for each plant, first bending the plant along the same axis as the touch treatment, and then a second test perpendicular to this axis. Plants from both treatments that had obvious poor establishment or visible breakage of nodal roots were excluded.

### Texture analysis

Resistance of individual root segments to compressive force was measured using a TA-XT Plus Texture Analyzer (Texture Technologies), equipped with a 55 mm, flat compression probe. Compression measurements were carried out using force target mode (25000 mN) and return to start mode with a probe speed of 0.2 mm/sec. Excised nodal roots were cut into 1.5 cm segments and placed flat on the stage of the texture analyzer with compressive force applied laterally relative to the root’s original orientation in soil. For each test, the force required to penetrate the sample 25%, 50%, and 75% of its diameter was calculated. First, the diameter of the segment was calculated by identifying the point where the probe contacted the root (the position at which the measured force first exceeded 2.5 mN) and subtracting that from the probe starting position. Force displacement curves were normalized to root diameter. To calculate displacement at a given percentage, normalized force values were averaged from the interval of n-1 to n+1, i.e., force at 25% represents the average of values from 24% to 26%.

### RNA sequencing and analysis

To harvest root tissue for RNA extraction a longitudinal cut was made in the pot. This allowed for plant removal from the pot without an uprooting force on the plant that might confound responses to mechanical treatment. Leaf nodal roots were quickly excised at the shoot root junction, and further dissected into a proximal basal region (0-1 cm from the shoot-root junction) and a distal basal region (3-4 cm), excluding the 1 cm interval between these regions to ensure a representative signal from thickened and unthickened regions. Segments of three separate roots per plant were pooled. Total RNA was extracted with the Plant RNeasy mini kit (Qiagen) according to manufacturer instructions with on-column treatment of RNase-free DNase I. RNA quality was evaluated with a Nanodrop spectrophotometer 2000c and Qubit 2.0 fluorometer. RNA sample quality, library preparation, and RNA sequencing were performed under contract by GeneWIZ. Quality of sequencing data was assessed with FastQC v0.11.9 (Andrews, 2010) and low-quality reads were trimmed using TrimGalore v0.6.10. High quality reads were mapped to the Bd21 genome v3.2 with Hisat2 v2.2.1 (Kim et al., 2015). Per-gene reads were quantified using StringTie v1.3.4 (Pertea et al., 2015), and read counts were normalized and assessed for differential expression using the DESEQ2 R package (Love et al., 2014). Gene ontology enrichment analysis was carried out with KOBAS v3.0 (Bu et al., 2021). In a preliminary analysis, principal component analysis showed that one pair of control distal and control proximal samples from the same plant clustered away from other replicates. These samples were eliminated from analysis and differentially expressed genes and enriched GO terms were recalculated. To find groups of genes with similar expression patterns, we first filtered to include differentially expressed genes. Transcript abundance was then standardized by computing z-scores within each treatment. The distance between each pair of genes was calculated based on the euclidean method and hierarchically clustered according to the complete linkage method. *Cis*-regulatory sequence analysis of differentially expressed genes was conducted as previously described using both a *de novo* k-mer approach and testing for overrepresented motifs present in the HOMER v4.10 *A. thaliana* DAP-seq database of 2 kb of sequence upstream of the transcriptional start site using Hypergeometric Optimization of Motif EnRichment suite (Heinz et al., 2010; Moore et al., 2022; Coomey et al., 2023). Raw read data was deposited in the European Nucleotide Archive for public access (Accession PRJEB71480).

### DNA affinity purification sequencing

DNA affinity purification was carried out as previously described (Coomey et al., 2023). DAP-seq data were first trimmed using BBtools bbduk. To identify binding target regions, reads were mapped to the Bd21 genome v3.2 using bowtie2 and peak calling was done using MACS3 (Zhang et al., 2008; Langmead and Salzberg, 2012). Motif calling was performed using Meme-suite v5.5.5 and the +/- 30 bp of the 100 tallest peaks (Bailey et al., 2015). We considered peaks containing the most significant motif and assigned those to genes using the ChIPseeker package in R (Yu et al., 2015). Raw read data was deposited in the European Nucleotide Archive for public access (Accession PRJEB71398).

### Statistical analysis

Histological and gross phenotyping experiments were analyzed using analysis of variance (ANOVA). Pairwise differences were then assessed using Tukey’s HSD test with a *p* < 0.05 cutoff. In RNA-seq analyses, adjusted *p* values were calculated using a Benjamini-Hochberg test. Figures were created with BioRender.

## Accession numbers

4CL2 (Bradi3g05750), C3’H1 (Bradi2g21300), C3H (Bradi1g65820), C4H1 (Bradi2g53470), CAD1 (Bradi3g06480), CCoAOMT2 (Bradi3g39420), CCR1 (Bradi3g36887), CESA1 (Bradi2g34240), CESA2 (Bradi1g04597), CESA3 (Bradi1g54250), CESA4 (Bradi3g28350), CESA5 (Bradi1g29060), CESA6 (Bradi1g53207), CESA7 (Bradi4g30540), CESA8 (Bradi2g49912), CESA9 (Bradi1g02510), CESA10 (Bradi1g36740), CSLF6a (Bradi3g16307), COBL (Bradi1g59880), COMT6 (Bradi3g16530), F5H (Bradi3g30590), GT43A (Bradi5g24290), GT47D3 (Bradi2g59400), GT61 (Bradi2g01480), HCT1 (Bradi5g14720), HCT2 (Bradi3g48530), LAC5 (Bradi1g66720), LAC6 (Bradi1g74320), LAC8 (Bradi2g23370), LAC12 (Bradi2g54740), MYB1 (Bradi4g06317), MYB2 (Bradi1g10470), MYB36 (Bradi23710), MYB41 (Bradi2g36730), MYB104 (Bradi5g20130), MYB69 (Bradi3g42430), PAL2 (Bradi3g49260), PMT (Bradi2g36910), PTAL1 (Bradi3g49250), SLR1 (Bradi1g11090), SND2R (Bradi2g462197), SWAM1 (Bradi2g47590), SWAM3 (Bradi2g40620), SWN2 (Bradi5g16917), SWN3 (Bradi3g50067), SWN4 (Bradi1g52187), SWN5 (Bradi5g27467), SWN6 (Bradi3g13117), SWN7 (Bradi1g50057), SWN8 (Bradi3g13727), XTH8 (Bradi3g18600).

## Acknowledgments

We thank H. Kim, G. Moreira Lana, A. Crosby for assistance with the texture analyzer and J. Chambers for assistance with microscopy.

## Author contributions

IWM, SPH, RCO designed the research; IWM, CFP, LTA, ELF, GAG, YZ performed research; LAB, RCO contributed new reagents; IWM, BK, LAB analyzed data; and IWM, SPH wrote the paper.

## Supplementary Data

**Supplementary Figure 1. *Brachypodium distachyon* nodal root anatomy.** Ep: Epidermis, Ex: Exodermis, CF1: Cortex File 1, CF2: Cortex File 2, CF3: Cortex File 3, P: Pith,Imx: Inner Metaxylem, Omx: Outer Metaxylem, Cs: Casparian Strip. Micrograph of a nodal root stained with phloroglucinol-HCl. Scale bar indicates 100 μm.

**Supplementary Figure 2. Quantification of secondary wall development as a function of root length.** Cell wall thickness **(A)** and phloroglucinol-HCl stain intensity (8-bit grayscale saturation value) **(B)**. Each point represents the cell type mean for one root. R and *p*-values reflect Pearson’s correlation.

**Supplementary Figure 3. Comparison of lignin and suberin localization.** Transverse sections of the proximal basal region of the same nodal root, stained with Sudan Black B for suberin **(A)** and phloroglucinol-HCl for lignin **(B)**. Scale bars indicate 100 μm.

**Supplementary Figure 4. Gross phenotypes of control and mechanically stimulated plants. (A).** Representative control (left 3 plants) and treated (right 3 plants) plants. Scale bar indicates 5 cm. Quantification of plant height **(B)**. Student’s t-test, ****p ≤ 0.0001.

**Supplementary Figure 5. Distal root phenotypes of hydroponically grown roots.** Representative transverse sections of the distal region (∼4 cm from the shoot-root junction) of plants grown under control **(A)** and 10 µM GA3 **(B)** treatment.

**Supplementary Figure 6. Analysis of gene expression by RNA-seq of nodal roots.** Principal component analysis plot illustrating the relationship among RNA-seq samples **(A).** Venn diagram of differential expressed transcripts between tissue and treatment comparisons **(B).**

**Supplementary Figure 7. Gene Ontology (GO) analysis depicting the functional categorization of differentially expressed genes among nodal root samples.** The analysis depicts enriched biological processes. GO terms enriched among stringently differentially expressed transcripts (*p* ≤ 0.01) in each comparison set.

**Supplementary Figure 8. Cis-regulatory sequences enriched among hierarchical clusters of differentially expressed genes in nodal roots.** *motif previously reported as enriched among genes that show a transcriptional response to mechanical stimulus (Coomey et al. 2023, Moore et al. 2022).

**Supplementary Figure 9. Transcript abundance measured by RNA-seq of selected secondary cell wall regulatory transcription factor genes in nodal root samples.** *SECONDARY WALL NAC* **(A)** and secondary wall associated MYB **(B)** family transcription factors. ∗ adj-*p* ≤ 0.05, ∗∗ adj-*p* ≤ 0.01, ∗∗∗ adj-*p* ≤ 0.001, ∗∗∗∗ adj-*p* ≤ 0.0001. N = 5 control and 4 brushed plants.

**Supplementary Figure 10. Transcript abundance measured by RNA-seq of cellulose, hemicellulose, and lignin biosynthesis associated genes in nodal root samples.** Expression of *CELLULOSE SYNTHASE A* (*CESA*) family **(A)**, lignin biosynthesis pathway **(B)**, and hemicellulose associated glycosyltransferase **(C)** genes. ∗ adj-*p* ≤ 0.05, ∗∗ adj-*p* ≤ 0.01, ∗∗∗ adj-*p* ≤ 0.001, ∗∗∗∗ adj-*p* ≤ 0.0001. N = 5 control and 4 brushed plants.

**Supplementary Table 1.** Genes differentially expressed between control and mechanically stimulated proximal leaf nodal roots.

**Supplementary Table 2.** Genes differentially expressed between proximal control and distal control leaf nodal roots.

**Supplementary Table 3.** GO term analysis of genes differentially expressed in the proximal root samples relative to the distal root. Category codes, F= molecular function, C = cellular component, P = biological process.

**Supplementary Table 4.** Hierarchical cluster analysis of differentially expressed genes using the complete linkage method.

**Supplementary Table 5.** All putative cis-regulatory elements enriched among differentially expressed genes at each cluster identified using the k-mer approach.

**Supplementary Table 6.** DNA affinity purification sequencing of SECONDARY WALL NAC7 (Bradi1g50057).

## DATA AVAILABILITY STATEMENT

The data that support the findings of this study are openly available at PRJEB71398 and PRJEB71480 in the European Nucleotide Archive and https://hazenlab-umass.shinyapps.io/LNR-RNAseq/.

