## Supplemental Figures 1-10 for "Shoring up the base: the development and regulation of cortical sclerenchyma in grass nodal roots"

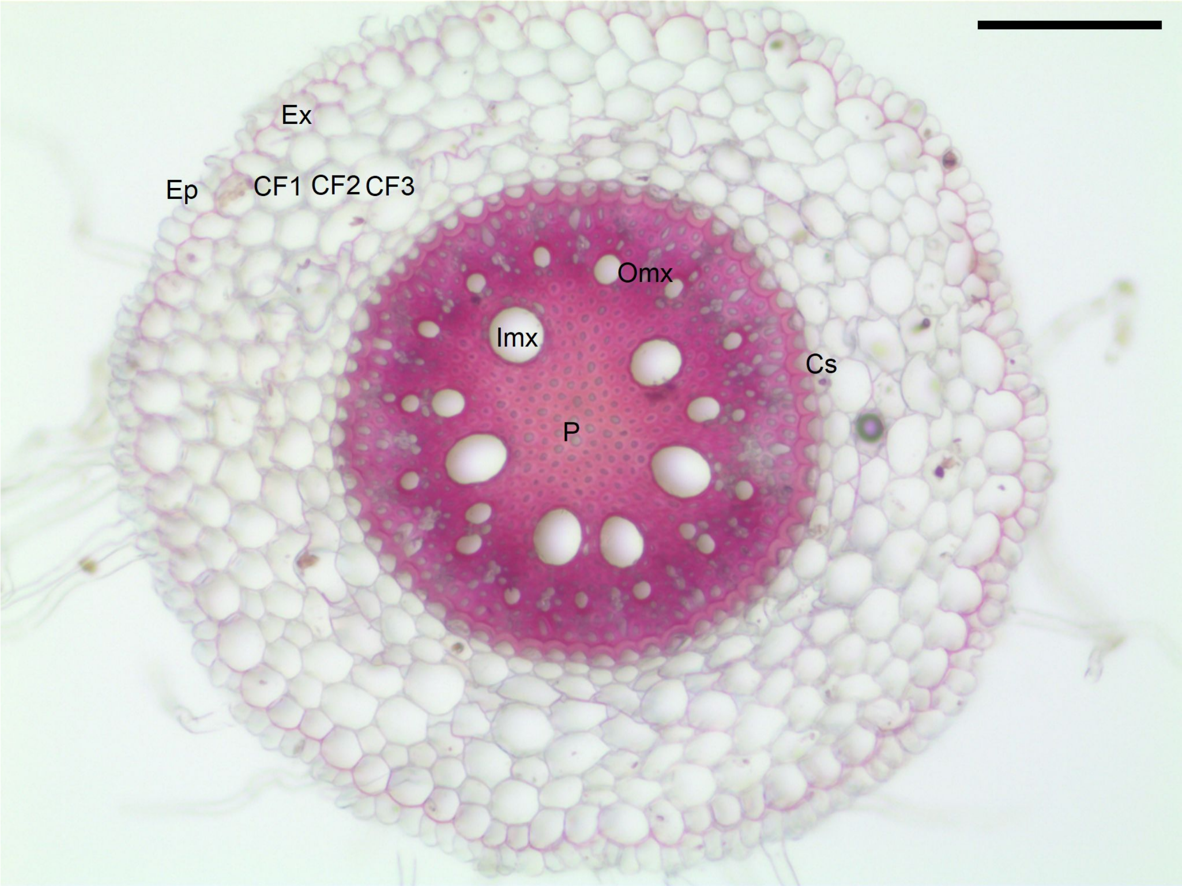


**Supplementary Figure 1. *Brachypodium distachyon* nodal root anatomy.** Ep: Epidermis, Ex: Exodermis, CF1: Cortex File 1, CF2: Cortex File 2, CF3: Cortex File 3, P: Pith,Imx: Inner Metaxylem, Omx: Outer Metaxylem, Cs: Casparian Strip. Micrograph of a nodal root stained with phloroglucinol-HCl. Scale bar indicates 100 μm.

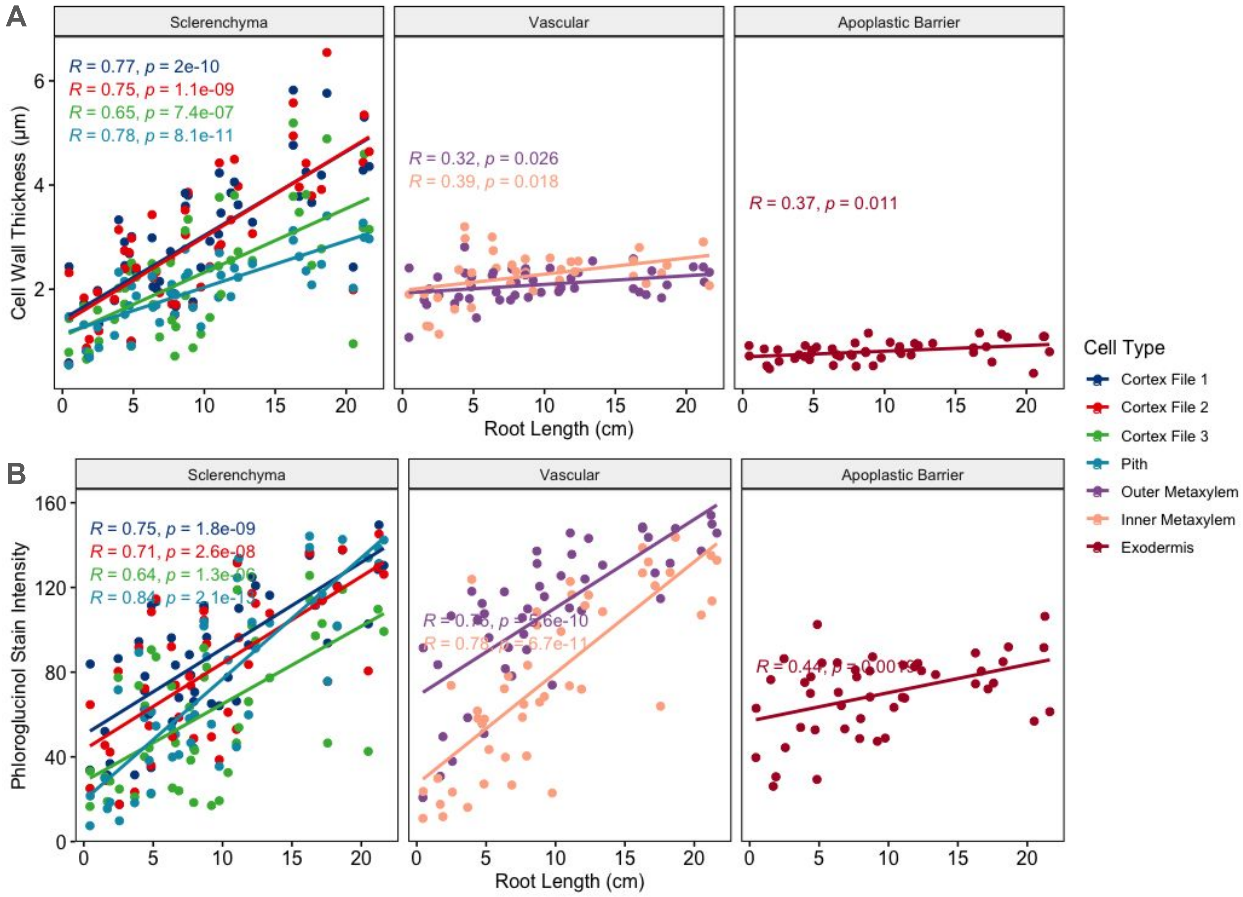


**Supplementary Figure 2. Quantification of secondary wall development as a function of root length.** Cell wall thickness **(A)** and **p**hloroglucinol-HCl stain intensity (8-bit grayscale saturation value) **(B)**. Each point represents the cell type mean for one root. R and *p*-values reflect Pearson's correlation.

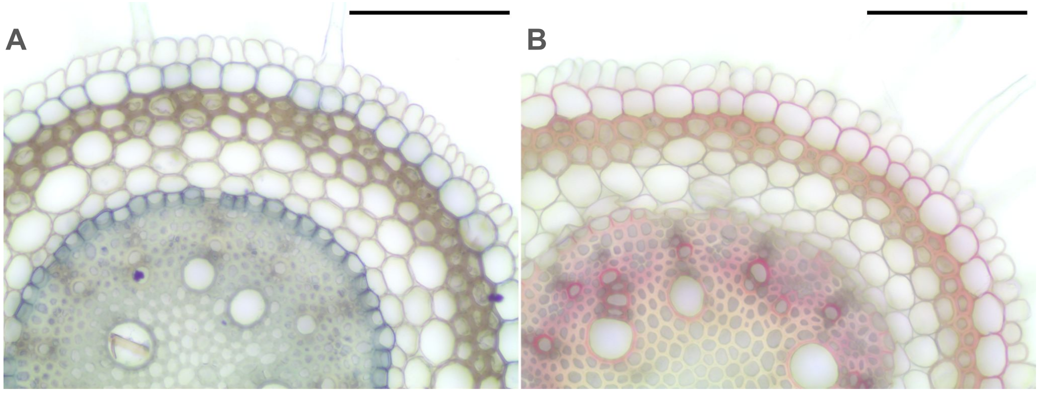


**Supplementary Figure 3. Comparison of lignin and suberin localization.** Transverse sections of the proximal basal region of the same nodal root, stained with Sudan Black B for suberin **(A)** and phloroglucinol-HCl for lignin **(B)**. Scale bars indicate 100 µm.

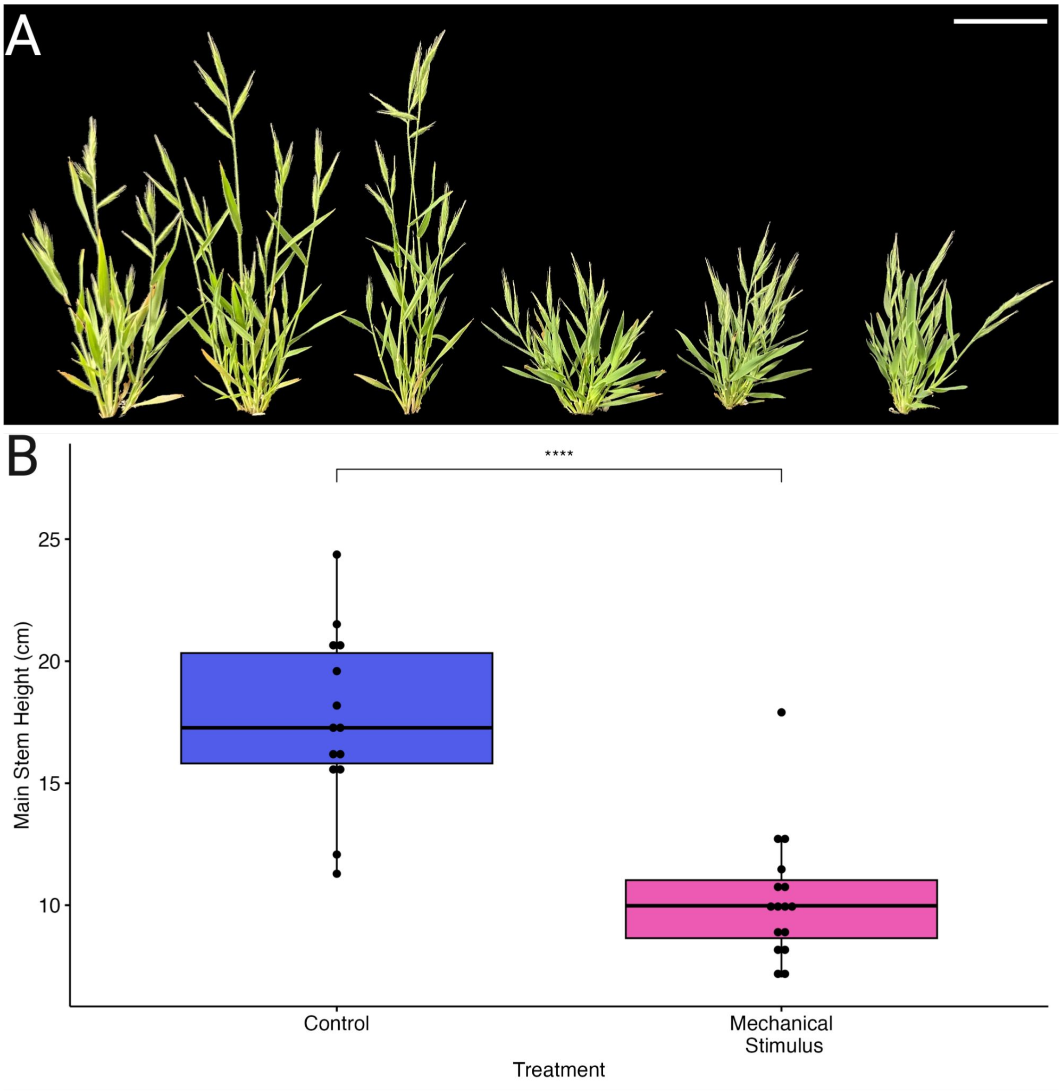


**Supplementary Figure 4. Gross phenotypes of control and mechanically stimulated plants. (A).** Representative control (left 3 plants) and treated (right 3 plants) plants. Scale bar indicates 5 cm. Quantification of plant height **(B)**. Student's t-test, ****$p \leq 0.0001$.

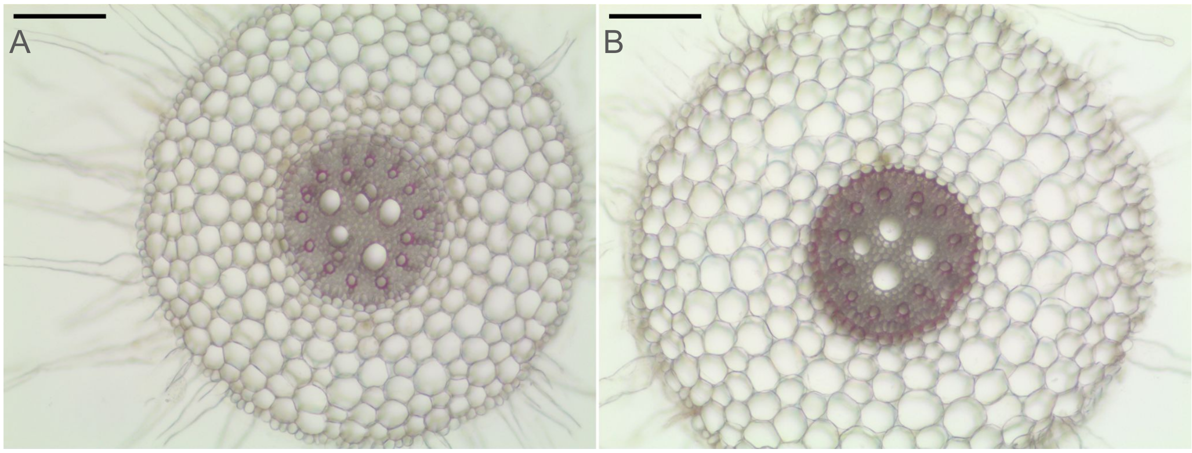


**Supplementary Figure 5. Distal root phenotypes of hydroponically grown roots.** Representative transverse sections of the distal region (~4 cm from the shoot-root junction) of plants grown under control **(A)** and 10 µM GA3 **(B)** treatment**.**

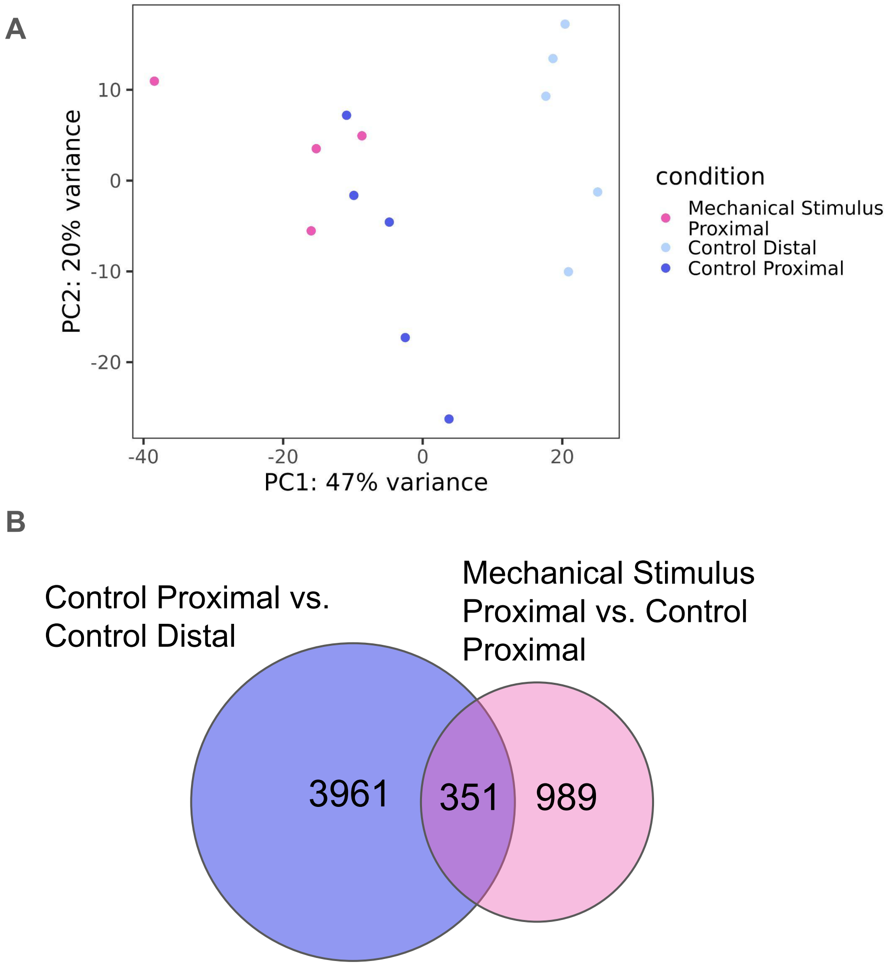


**Supplementary Figure 6. Analysis of gene expression by RNA-seq of nodal roots.** Principal component analysis plot illustrating the relationship among RNA-seq samples **(A)**. Venn diagram of differential expressed transcripts between tissue and treatment comparisons **(B)**.

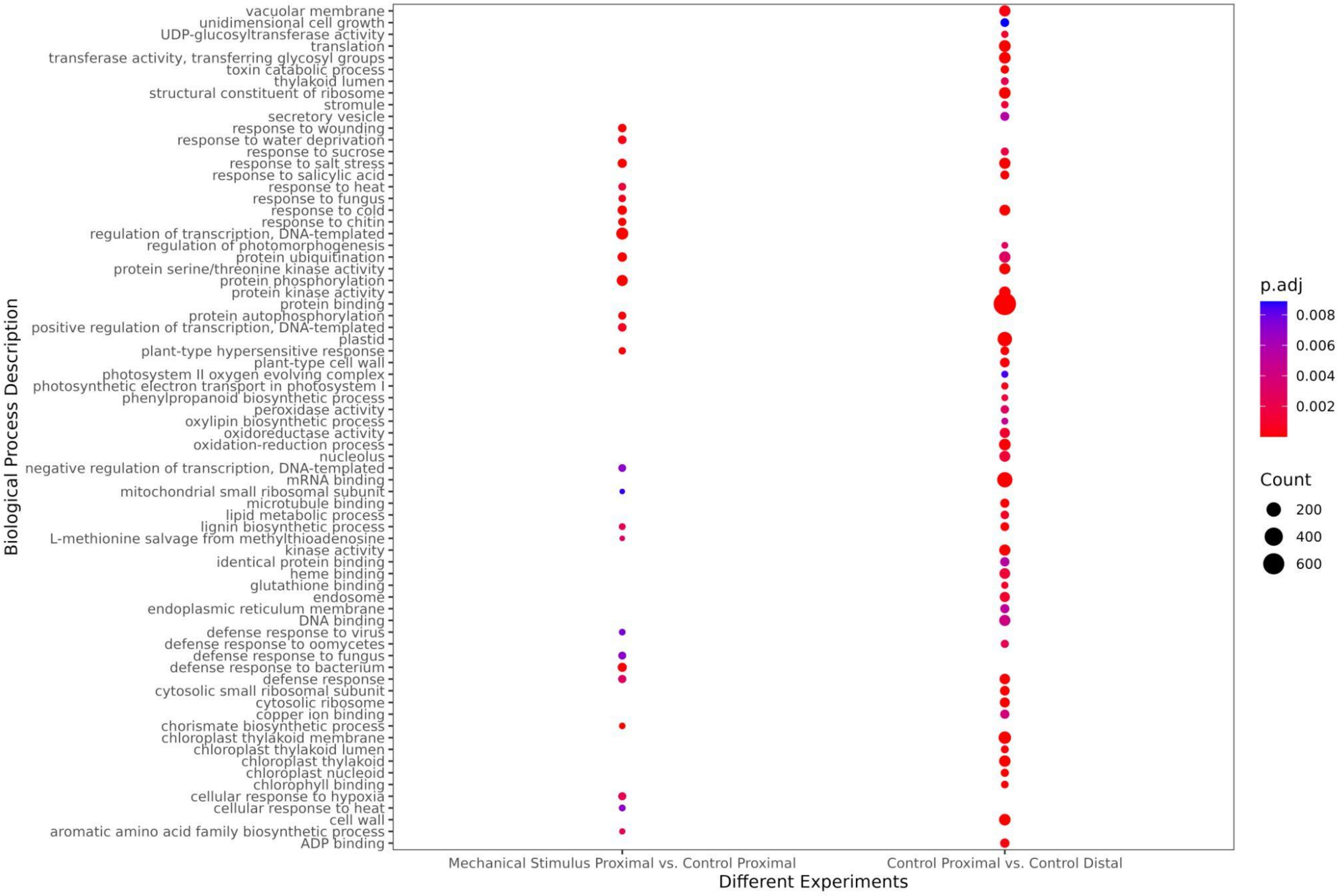


**Supplementary Figure 7. Gene Ontology (GO) analysis depicting the functional categorization of differentially expressed genes among nodal root samples.** The analysis depicts enriched biological processes. GO terms enriched among stringently differentially expressed transcripts ($p \leq 0.01$) in each comparison set.

**Overrepresented *cis*-regulatory sequences**

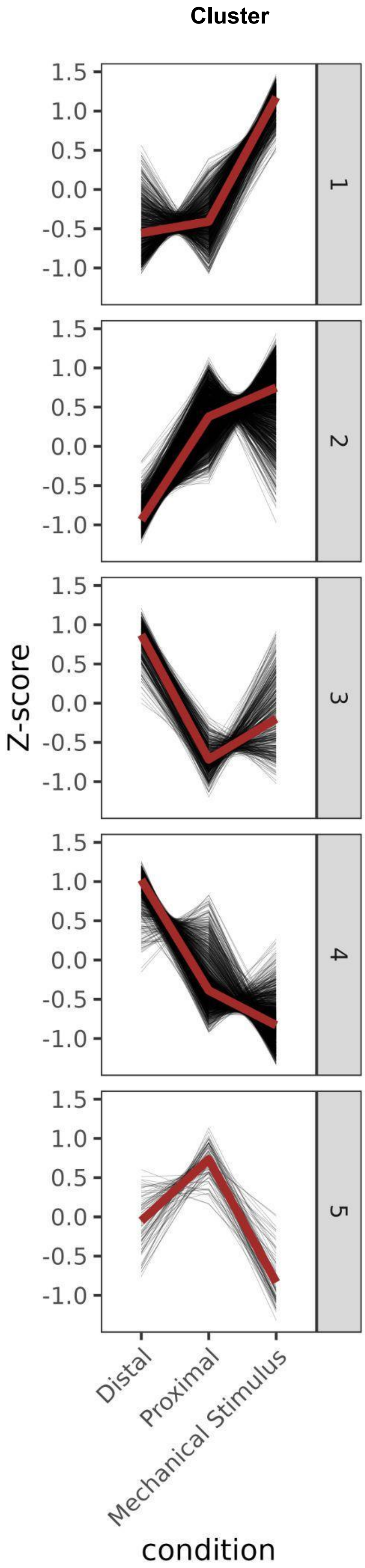


Cluster 1:
ACC(A/T)ACC (AC element)*
ACGCGC (FAR1 binding site)*
CACGCG (G/A-box)*
C(C/T)TTTT (P-box)*
CGCGTT (RSRE)*

Cluster 2:
AAATATCT (MYB-related binding site)
(A/G)CCGAC (AP2-like binding site)*
AGATATTT (ND binding site)
ACGTGGC (ABRE)
CTTT(T/A) (P-box)*
CACGTG (G-box)*
G(C/A)CACGT (G-box)*
GANNNGA(C/T) (Homeobox binding site)
GATAA (MYB-related binding site)
GGNCCCAC (Site II element)*

Cluster 3:
CGCGTT (RSRE)*
TTGAC(C/T) (W-box)*

Cluster 4:
AATCAT (Homeobox binding site)
CTTTTT (P-box)*

Cluster 5:
CCTAAA (MYB-related binding site)

**Supplementary Figure 8. Cis-regulatory sequences enriched among hierarchical clusters of differentially expressed genes in nodal roots.** *motif previously reported as enriched among genes that show a transcriptional response to mechanical stimulus (Coomey et al. 2023, Moore et al. 2022).

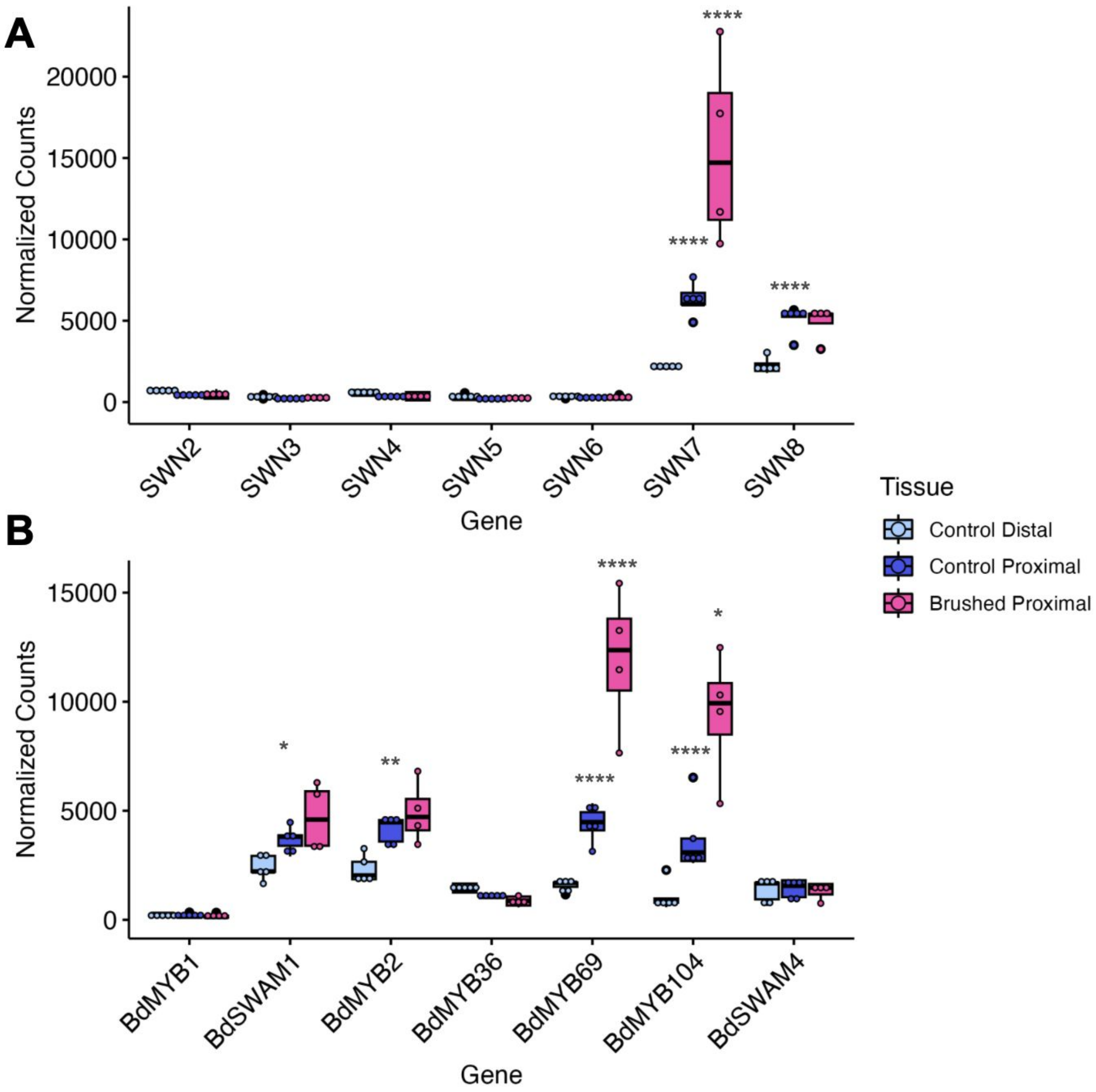


**Supplementary Figure 9.** Transcript abundance measured by RNA-seq of selected secondary cell wall regulatory transcription factor genes in nodal root samples. *SECONDARY WALL NAC* **(A)** and secondary wall associated MYB **(B)** family transcription factors . ∗ adj-$p \leq 0.05$, ∗∗ adj-$p \leq 0.01$, ∗∗∗ adj-$p \leq 0.001$, ∗∗∗∗ adj-$p \leq 0.0001$. N = 5 control and 4 brushed plants.

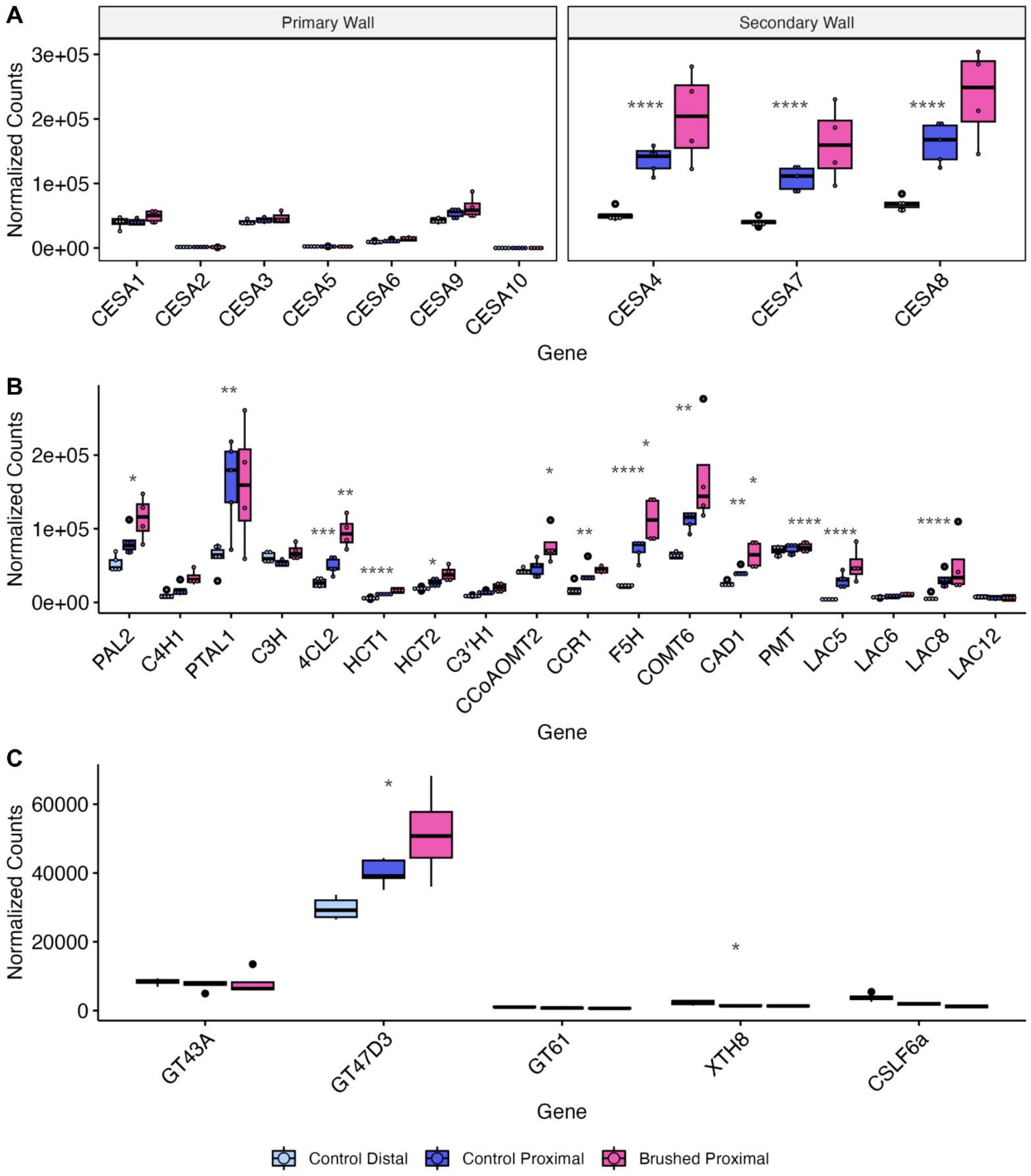


**Supplementary Figure 10. Transcript abundance measured by RNA-seq of cellulose, hemicellulose, and lignin biosynthesis associated genes in nodal root samples.** Expression of *CELLULOSE SYNTHASE A* (*CESA*) family **(A),** lignin biosynthesis pathway **(B)**, and hemicellulose associated glycosyltransferase **(C)** genes. ∗ adj-$p \leq 0.05$, ∗∗ adj-$p \leq 0.01$, ∗∗∗ adj-$p \leq 0.001$, ∗∗∗∗ adj-$p \leq 0.0001$. N = 5 control and 4 brushed plants.
